# A cognitive map of subjective value space for human risky choice

**DOI:** 10.64898/2026.05.19.726239

**Authors:** Mark A. Orloff, Seongmin A. Park, Jake Blumwald, Philippe Domenech, Erie D. Boorman

## Abstract

Individuals are thought to make choices based on subjective valuations of options that integrate multiple attributes into a unified subjective value signal in the brain’s “value coding system.” How these attributes are transformed into a subjective valuation remains poorly understood. One candidate is the “cognitive mapping system,” which efficiently represents relational, multi-dimensional information and has been proposed to afford novel inferences. We develop a risky decision-making task and use fMRI to show that a two-dimensional (2D) subjective value space of reward probability and amount is represented in the cognitive mapping system as both a grid-like representation of decision vectors and a 2D “positional” code in entorhinal cortex (EC) and medial prefrontal cortex (mPFC). The strength of grid-like and subjective value representations are positively correlated, suggesting the cognitive mapping and value coding systems work in tandem. These findings bridge these systems and support a new framework for how the brain constructs values.

## Introduction

A predominant view in decision neurosciences holds that reward value is represented in the brain as a unified, one-dimensional signal in “value coding regions.” Value-related modulation of neural activity has been reported in ventromedial prefrontal cortex (vmPFC), orbitofrontal cortex, posterior cingulate cortex (PCC), ventral striatum, basolateral amygdala, dorsomedial frontal cortex (dmFC), and dorsolateral prefrontal cortex (dlPFC), among other regions ^1,2^. Many of these value signals reflect the weighted integration of attributes, such as reward probability, amount, effort, and delay, of different rewards, such as money, juice, or social rewards, in a “common neural currency” ^3,4^. Crucially, these signals reflect the *subjective* value of rewards—how much a particular outcome is worth based on an individual’s preferences. Despite the ubiquity of such value signals, the mechanisms by which subjective value is constructed from an option’s attributes remain unknown.

One potential mechanism by which individuals may perform such value construction under some circumstances is via the brain’s cognitive mapping system. Cognitive maps are thought to arise, in part, from so-called ‘grid cells’ in the entorhinal cortex (EC), and, at least in primates, medial prefrontal cortex (mPFC) ^5–7^. This neuronal population representation efficiently codes relational information and has been proposed to enable flexible inference and generalization ^8–12^. Map-like representations in these regions were first identified through physical navigation tasks in both animal models and humans ^5–7,13^, but more recently have been identified in “abstract” relational tasks—non-spatial tasks with 2D relational structures even when they are never explicitly revealed to the participant but entail “traversing” the 2D space ^12–21^. In the context of value-based decision-making, this cognitive mapping system could aid in many everyday situations where the values of new options are unknown, inferences based on past experiences can be generalized, and value needs to be constructed on the fly. For example, divorcing the sensory attributes of particular options (e.g., the redness and texture of a berry) from an option’s general value (as represented as a common neural currency) allows for flexible compositional generalization to construct the value of new options (e.g. a new berry).

Recent work in primates has only begun to examine the relationship between cognitive maps and value based decision-making, producing conflicting results. One fMRI study in humans found no evidence of a grid-like representation during inter-temporal choices but did identify subjective value signals in the brain’s value coding system, and suggested the brain only uses a value coding system for value-based choices and a grid coding system for a distinct set of relational tasks ^22^. More recently, however, grid-like representations have been identified in monkey vmPFC using fMRI and direct electrophysiological recordings in value-based decision-making tasks ^23,24^, although, importantly, such coding has not been reported in the EC and was not linked to subjective preferences. Moreover, monkey hippocampal neurons were shown to represent a monkey’s “position” and trajectory when learning a three-dimensional reward value space, consistent with so-called “place cells” for an abstract value space ^25^. Whether humans likewise utilize a cognitive map to represent and infer over an abstract value space for value-based decisions, and whether this representation reflects subjective preferences is thus unclear and a topic of ongoing debate in the field.

In spatial and abstract navigation, other work suggests that spatial and non-spatial cognitive maps distort based on environmental characteristics ^26–28^. For example, grid-like and “position” coding representations distort based on an environment’s shape ^29^, the location of rewards ^30,31^, and task demands placed on inferences ^32^. In humans, these distortions are reflected in behavioral biases such as response time ^32^ and location precision ^33^. These intriguing findings raise the question of whether the cognitive mapping system also represents distorted value information according to subjective preferences in humans.

Here, we developed a novel risky decision-making task with two relevant decision attributes (reward probability and amount) to test whether humans represent an abstract value space as a cognitive map and infer decision vectors over the value space during value-based choice. We utilize computational modeling of risky choices to estimate individual nonlinear distortions in each value dimension to construct subjective value spaces for each participant. We hypothesized that we would observe two key components of the cognitive mapping system during risky decision-making: 1) grid-like modulation of inferred decision vectors, and 2) a generalized two-dimensional (2D) positional code of options in the value space in EC and mPFC. If the cognitive mapping system reflects individuals’ subjective value spaces, these features should be strongest after accounting for these distortions.

## Results

### Risky decision-making task

To create an abstract value space, we trained individuals (N = 35) to learn stimulus characteristics of two sets of shapes, each with two continuously varying dimensions corresponding to probability (10–100%) and amount ($1–10; Fig. 1B) of reward. Participants were trained piece-meal, such that they learned one dimension at a time. Crucially, participants never saw the 2D value space. Additionally, participants only saw a subset of all possible stimuli, such that only 32 of 88 (36%) shapes per set were seen before the main scanning session.

**Figure 1.**
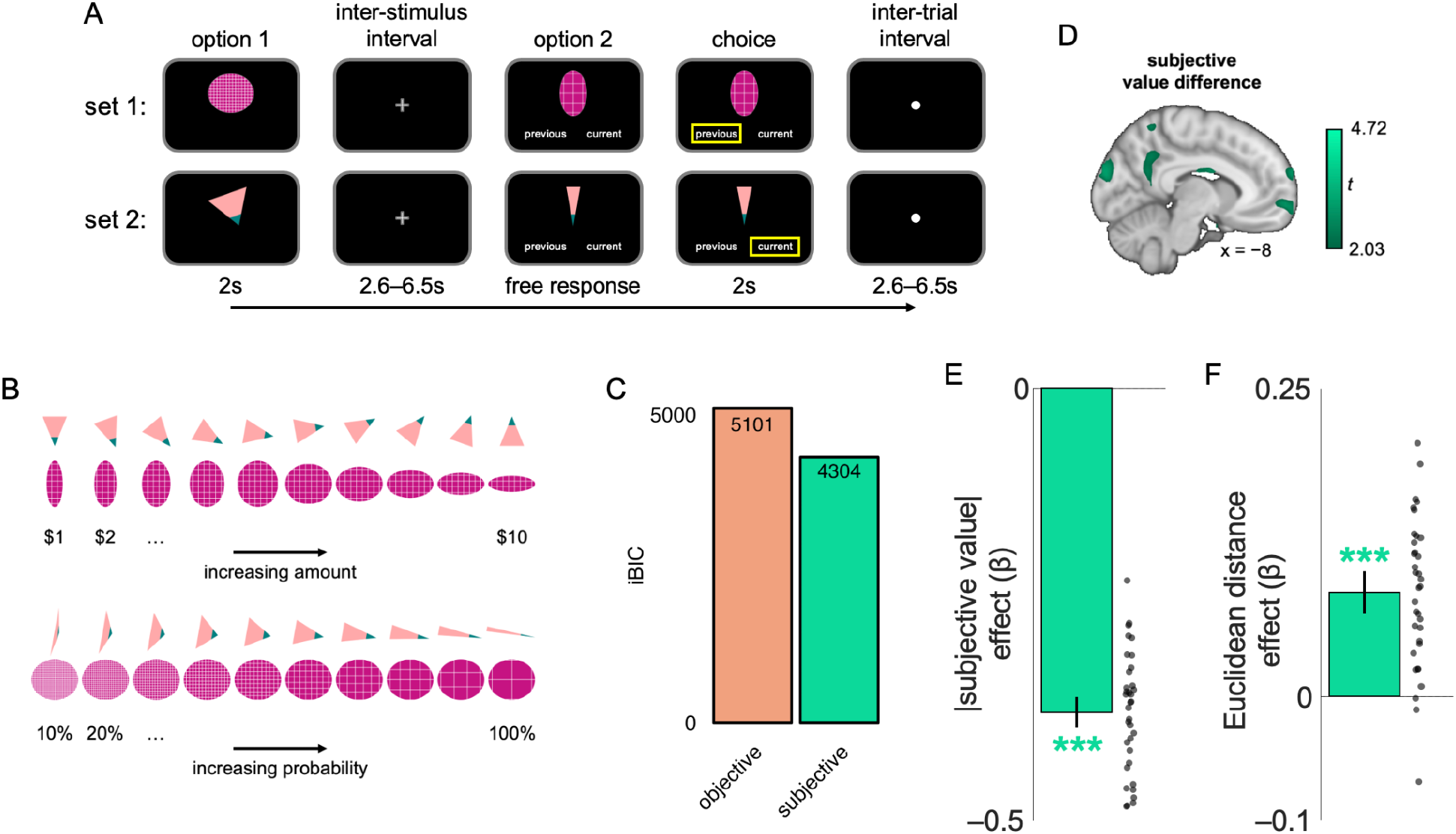
Task structure, model comparison, value coding regions, and response time regression estimates. **A)** In this task, participants are asked to make choices between sequentially shown shapes based on the amounts and probabilities associated with them while undergoing fMRI scanning. **B)** Participants learned about two sets of shapes, “triangles” and “ovals,” where each set had two feature dimensions, corresponding to amount and probability, along which they continuously varied. **C)** The subjective value computational model explained participants’ choices better than the objective value model. **D)** Whole-brain maps of the subjective value difference between the chosen and unchosen option at the time of the second shape onset. The map is thresholded at *P*_uncorrected_ < 0.025 for visualization purposes. **E)** A mixed effects regression shows that across subjects, as |subjective value difference| between shape options increases, RT decreases. **F)** A mixed effects regression on the residuals from **E** shows that across subjects, as Euclidean distance between shape options increases, RTs increase. Bar plots and error bars are the fixed effect and SEM of the estimate. The dots on are the reconstructed β values for each subject (i.e., the fixed effect β + random effects β for each subject). \*\*\**P* < 0.001

After training, participants completed a risky decision-making task using these shapes as stimuli while undergoing fMRI scanning. In this task, two shapes were shown sequentially (Fig. 1A) and participants were asked to choose which shape they preferred based on the probabilities and amounts associated with each shape. This task was incentive compatible such that after each of three runs, one of the chosen shapes was randomly selected, and the amount associated with that shape was awarded to the participant depending on the shape’s probability. A risky decision-making task was used because it represents a canonical value-based decision making task and non-linear distortions in probability and amount are well-described and studied extensively in the broader neuro- and behavioral economics literature (e.g., ^34,35^).

### Behavioral and neural evidence of subjective weighting in risky decision-making

We utilized cumulative prospect theory to account for individuals’ subjective valuation of options ^34^. Specifically, we estimated a “subjective value” model including a probability weighting parameter and amount sensitivity parameter to account for nonlinear weighting of probability and amount, respectively. We compared this model to an “objective value” model, which computed expected value according to expected value theory, assuming no nonlinear distortions in probability or amount, using integrated Bayesian information criterion (iBIC), a modified version of BIC that accounts for a posterior distribution of parameter estimates rather than point estimates (where lower estimates indicate a better model fit; ^36^). As expected, the subjective value model was a better fit (objective iBIC = 5101, subjective iBIC = 4304; Fig. 1C).

Next, we sought to confirm that the value coding regions typically observed in value-based decision-making tasks were indeed representing the one-dimensional subjective value difference of the chosen option relative to the value of the unchosen option ^37–42^. We estimated two GLMs per subject, one assuming an objective value function (objective GLM_value_; i.e., without linear distortions to probability or amount; Extended Data Fig. 1A) and another using the subjective value parameters estimated for each participant to account for nonlinear distortions in value dimensions (subjective GLM_value_; Fig. 1D). We observed significant positive parameter estimates across both objective and subjective value in vmPFC (objective *P*_threshold-free cluster enhancement, small volume corrected (TFCE, SVC)_ < 0.05, subjective *P*_TFCE, SVC_ < 0.05, rostral vmPFC ROI), marginally in PCC (objective: *P*_TFCE, SVC_ = 0.059, subjective: *P*_TFCE, SVC_ = 0.051, vPCC), and significant negative parameter estimates in bilateral anterior insula (objective and subjective *P*s_TFCE_ < 0.05), bilateral dlPFC (objective and subjective *P*s_TFCE_ < 0.05), and dmFC (objective and subjective *P*s_TFCE_ < 0.05). Neural effects of risk difference are also consistent with prior risky decision-making tasks ^43–45^ (Extended Data Fig. 1B). Thus, in this novel task, individuals utilize similar value- and risk-sensitive regions as in other value-based and risky decision-making tasks.

### A cognitive map of value space

We hypothesized that a 2D value space would be represented as an abstract 2D cognitive map in the EC and mPFC. In this framework, each decision would utilize a simulated vector between the first and second presented shapes (i.e., a decision vector; Fig. 2A). Each decision vector is defined by a distance and angle through the value space. To test our main hypothesis, we further estimate subjective value spaces for each participant. To do so, we calculate new probability and amount axes according to individually estimated probability weighting and amount sensitivity parameters from choice behavior (normalized [0 1]). This new space will reflect nonlinear distortions in probability and amount, according to each individual’s unique estimates based on their subjective preferences (Fig. 2B).

**Figure 2.**
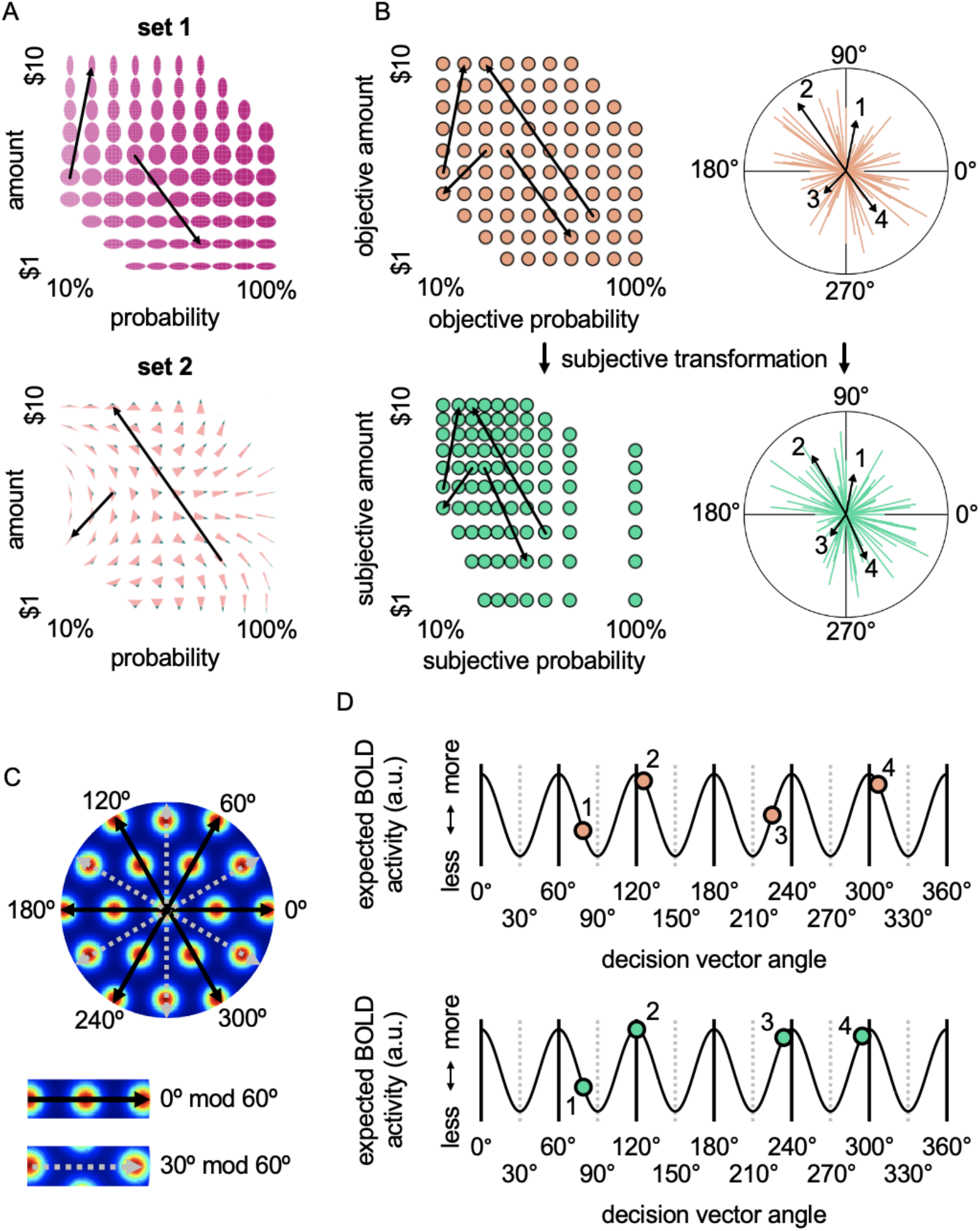
Decision vectors in objectively and subjectively defined value spaces. **A)** The full abstract value spaces for set 1 (top) and set 2 (bottom). Two example decision vectors are shown as arrows in each shape set. Each decision vector represents a choice in the 2D space where the start is the first shape shown and the end is the second shape. **B)** The example decision vectors are overlaid in a shared objective value space (top left), where each dot represents a shape. For every choice, a decision vector is obtained and the distance and angle through the abstract space is calculated (top right). For every individual, a transformation is applied to the space to account for their probability weighting and amount sensitivity. This results in a unique subjective value space for each subject (bottom left, one example subject). New decision vectors are calculated, and the angles and distances through the subjective space are displayed (bottom right). Note that the arrows in each plot in B correspond to the same four decision vectors in A. See Extended Data Fig. 3 to see subjective spaces and decision vectors for all subjects (subject 5 is shown here). **C)** The firing field of one grid cell is shown, where the hotter colors represent a greater firing rate as an individual moves through that location in the 2D space. This hexagonal “grid-like” activity was initially proposed to arise because of two key properties of grid cells ^13^: 1) within a given “module” grid cells all tend to be sensitive to the same movement directions (i.e., are aligned) ^47^, and 2) some “conjunctive” grid cells also encode head direction (i.e., are directionally sensitive) ^48^. Formal simulations confirm this as the most likely explanation for fMRI grid-like activity ^46^. Thus, movement at a particular angle, as measured across bulk activity of all grid cells in a particular voxel should produce a canonical hexagonal modulation. **D)** Because of these properties, for each angle of a decision vector in our task, we can make precise predictions about how much (relative) BOLD signal should be observed. Shown is the hypothesized relative BOLD activity for each of the four example decision vectors in **B** for the objective space (top) and the subjective space for one example subject (bottom). The numbers next to the dots in **D** correspond to the decision vectors shown in **A** and **B**. In our GLM_grid_, we specify a regressor that will reveal voxels with BOLD activity that correlates with this grid-like pattern.

### Behavioral evidence of a cognitive map in risky decision-making

After confirming our new task produces expected behavioral and neural findings consistent with other value-based decision-making tasks, we use mixed effects regression to analyze response times (RTs) to test if cognitive map representations are utilized in this task. We reasoned that if people are using cognitive maps in this task, RTs might vary with the distance between shapes for each decision vector, similarly to traveling in a physical space. Because distance between shapes is partially confounded with the value difference between shapes, we use a two-step procedure to account for value difference before testing for a relationship between distance and RT. First, we perform a mixed effects regression on the absolute subjective value difference (|U_diff_|, eq. 4). As expected, we observed that lower |subjective value difference| (i.e., greater decision difficulty) leads to slower RTs (β = −0.38, *P* < 0.001; Fig. 1E).

Next, we take the residuals of this model and further regress them against the Euclidean distance between the shapes in the abstract subjective value space (i.e., the length of the decision vector). This shows any effect of the length of the decision vector above and beyond the effect of subjective value difference on RT. Indeed, we observed a significant, albeit smaller, effect of Euclidean distance on RT (β = 0.085, *P* < 0.001; Fig. 1F), akin to physical navigation where longer distances take further to traverse. This effect is difficult to account for with current theories of value-based decision-making without appealing to a theory based on cognitive maps. Note that these results remain robust after accounting for other possible confounding task variables (Extended Data Fig. 2).

### Grid-like representation of an abstract value space

We hypothesized that individuals utilize a grid code to represent and infer vectors over an abstract value space. Using the calculated angles for each decision vector, we can predict the expected BOLD response for that particular decision vector based on the unique properties of grid cells’ firing fields, known as hexagonal symmetry ^13,46^ (Figs. 2C and 2D). Specifically, we test if BOLD response variance is better explained by hexagonal modulation with respect to the angles through the subjective space, separately defined for each individual. By design the angles for objective space, and for each individual’s subjective space, are well distributed throughout the unit circle (Kuiper’s test against a uniform distribution, all *P*s > 0.10), thus demonstrating a good sampling across decision vectors and sensitivity to detect hypothesized hexagonal modulation (see Extended Data Fig. 3 for all participants’ subjective value spaces and decision vectors). By design, these analyses are also unlikely to be confounded by differences in Euclidean distance because the differences between angles ‘on’ and ‘off’ grid are well matched across all subjects for objective and subjective spaces (all *P*s > 0.16).

**Figure 3.**
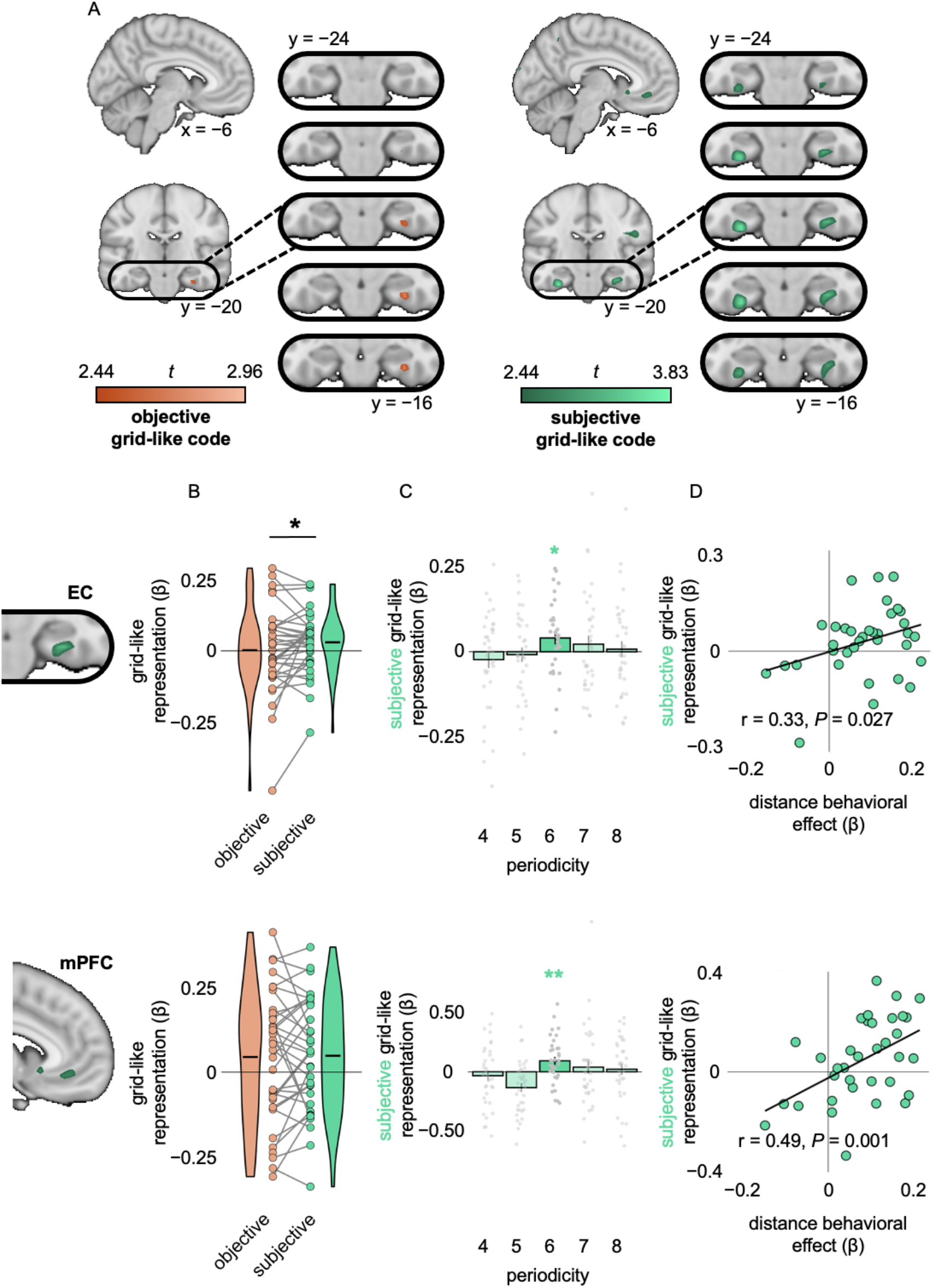
Grid-like representation of an abstract value space. **A)** Whole brain maps of grid-like representations of objective (left) and subjective (right) abstract value space at the time when the decision vector can be computed. These maps are thresholded at *P*_uncorrected_ < 0.01 for visualization. **B)** Extracted mean parameter estimates across subjects for grid-like representation in *a priori*, independently defined ROIs, right EC (top) and mPFC (bottom) for grid-like representation of objective (orange) and subjective (green) value space. In EC, but not mPFC, grid-like representation is stronger in the subjective value space. Grey lines indicate how individual subjects’ estimates change between value spaces. The black dot shows the group mean for each value space. **C)** A control analysis confirms that the grid-like representation in subjective value space is specific to 6-fold (rather than 4-, 5-, 7-, or 8-fold symmetry) in the EC (top) and mPFC (bottom). **D)** Grid-like representations are correlated with the distance behavioral effect across participants for both EC (top) and mPFC (bottom). Error bars represent standard error of the mean (s.e.m.). \**P* < 0.05, and \*\**P* < 0.01.

First, we test for a grid-like representation of decision vectors in the objective value space using a parametric analysis testing for hexagonal modulation (GLM_grid_). While some variance is explained in right EC at a reduced threshold, we did not observe any significant parameter estimates for hexagonal modulation in *a priori*, independently defined EC (right: *P*_TFCE, SVC_ = 0.40, left: *P*_TFCE, SVC_ = 0.30) or mPFC, after small volume correction (*P*_TFCE, SVC_ = 0.12; Fig. 3A). On the other hand, for decision vectors in the subjective value space, we observe significant parameter estimates in both right EC (*P*_TFCE, SVC_ < 0.05; peak MNI coordinates [x, y, z] = [30, −16, −36]) and mPFC (*P*_TFCE, SVC_ < 0.05; [−6, 36, −10]; Fig. 3A), but not left EC (*P*_TFCE, SVC_ = 0.21). Thus, grid-like coding was used to represent decision vectors through individuals’ subjective, but not objective value spaces, implying that the observed grid representation is distorted according to an individual’s subjective preference.

To formally test if the subjective grid-like representation is stronger than the objective grid-like representations, we extracted parameter estimates across anatomically defined ROIs. We observe larger estimates for the subjective grid-like code in right EC (*P* < 0.05, one-sided t-test), but similar estimates in mPFC (*P* = 0.43, one-sided t-test; Figs. 3B and 3C).

Control analyses confirm the robustness of our results of grid-like representation in a subjective value space. Significant parameter estimates were only observed in a 6-fold symmetry, as compared with 4-, 5-, 7-, and 8-fold control symmetries for right EC (6-fold: *P* < 0.05; 4-, 5-, 7-, and 8-fold *P*s > 0.19, one-sided t-test) and mPFC (6-fold: *P* < 0.05; 4-, 5-, 7-, and 8-fold *P*s > 0.25, one-sided t-test). Additionally, extracting parameter estimates across the twelve direction bins for visualization shows a similar pattern across ‘on’ and ‘off’ grid angles (Extended Data Fig. 4) across all angles sampled, showing that the overall effect was not systematically driven by a subset of angles. Finally, this result cannot be explained by a value or value difference effect, since the EC and mPFC subjective grid coding effect survives inclusion of subjective value difference regressors included in the same GLM (Extended Data Fig. 5). Because our task design deliberately included a majority of novel comparisons, theoretically benefiting from proposed computational advantages of using a cognitive map ^8,12^, we could further test whether this effect of hexagonal modulation was present when new decision vectors had to be constructed on the fly. Indeed, the EC and mPFC effects of hexagonal modulation remained consistent when only analyzing novel choices (i.e., excluding choices previously made in training; Extended Data Fig. 6). Together, these results demonstrate that the subjective grid-like representation in EC and mPFC is specific to the predicted hexagonal symmetry, is present across the full spectrum of decision vector angles, cannot be explained by value coding, and is present for novel choices. Further, this grid representation of decision vectors is strongest for the subjective, rather than objective, value space.

**Figure 4.**
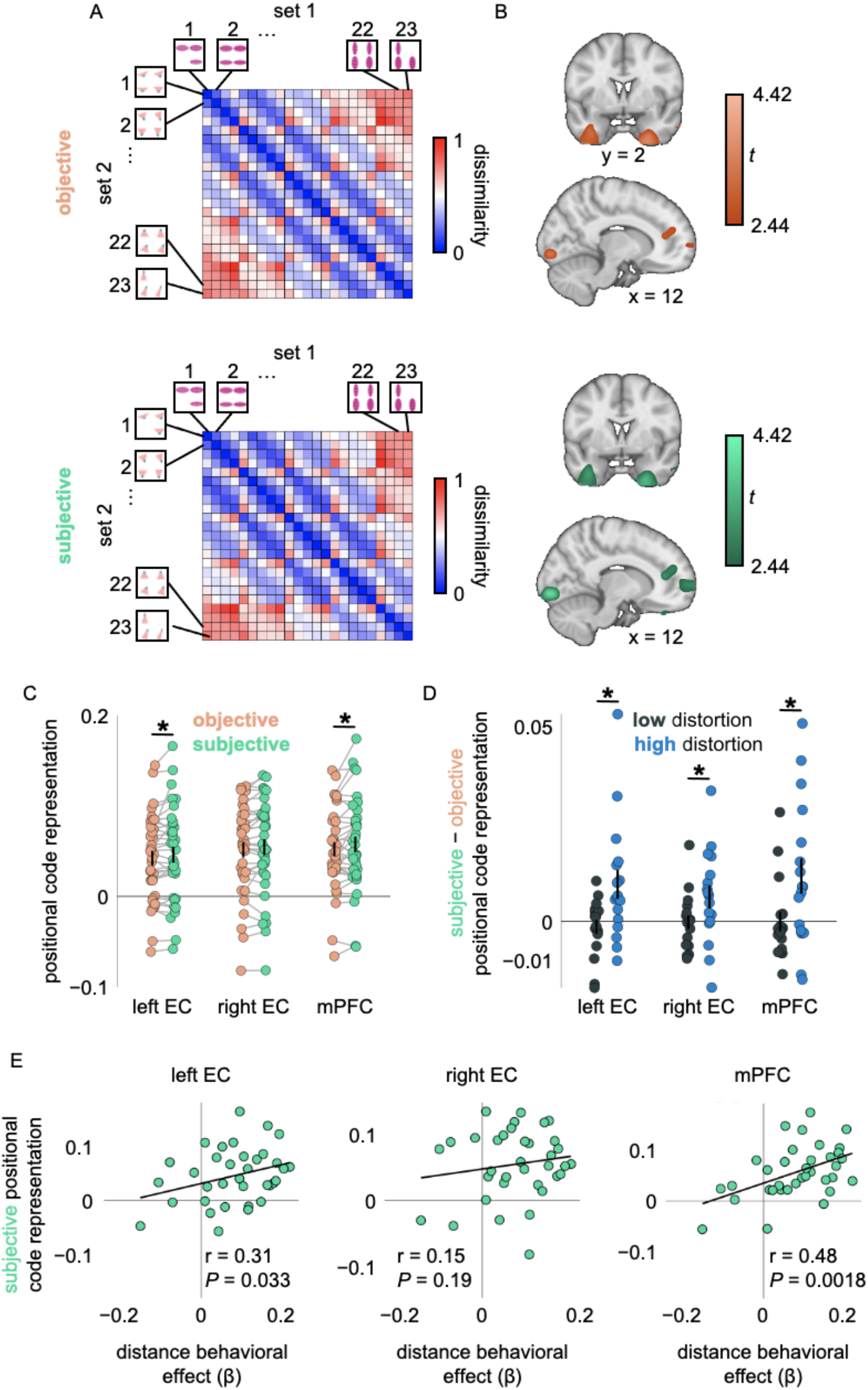
EC represents a positional code in a 2D subjective value space. **A)** The model RDM represents the (dis)similarity between each group of shapes. We test whether neural patterns in a whole-brain searchlight analysis resemble this dissimilarity pattern where shapes closer together in objective value space (top) and subjective value space (bottom, example subject) are represented more similarly than shapes that are further apart. **B)** Voxels across the whole-brain searchlight that demonstrate a 2D positional code in objective value space (top) and subjective value space (bottom). Displayed at *P*_uncorrected_ < 0.01. **C)** Positional code estimates across ROIs for objective and subjective distortion are shown. **D)** Subjects were placed into low and high distortion groups based on computational model parameter estimates. The difference between the subjective and objective RSA results in the top 33% of voxels are shown for each subject. Subjects with high distortion had significantly greater representation of a 2D subjective value space (relative to the objective value space) than the low distortion subjects. **E)** The subjective positional code representation is correlated with the distance behavioral effect across participants in left EC and mPFC. Error bars represent standard error of the mean (s.e.m.). \**P* < 0.05.

**Figure 5.**
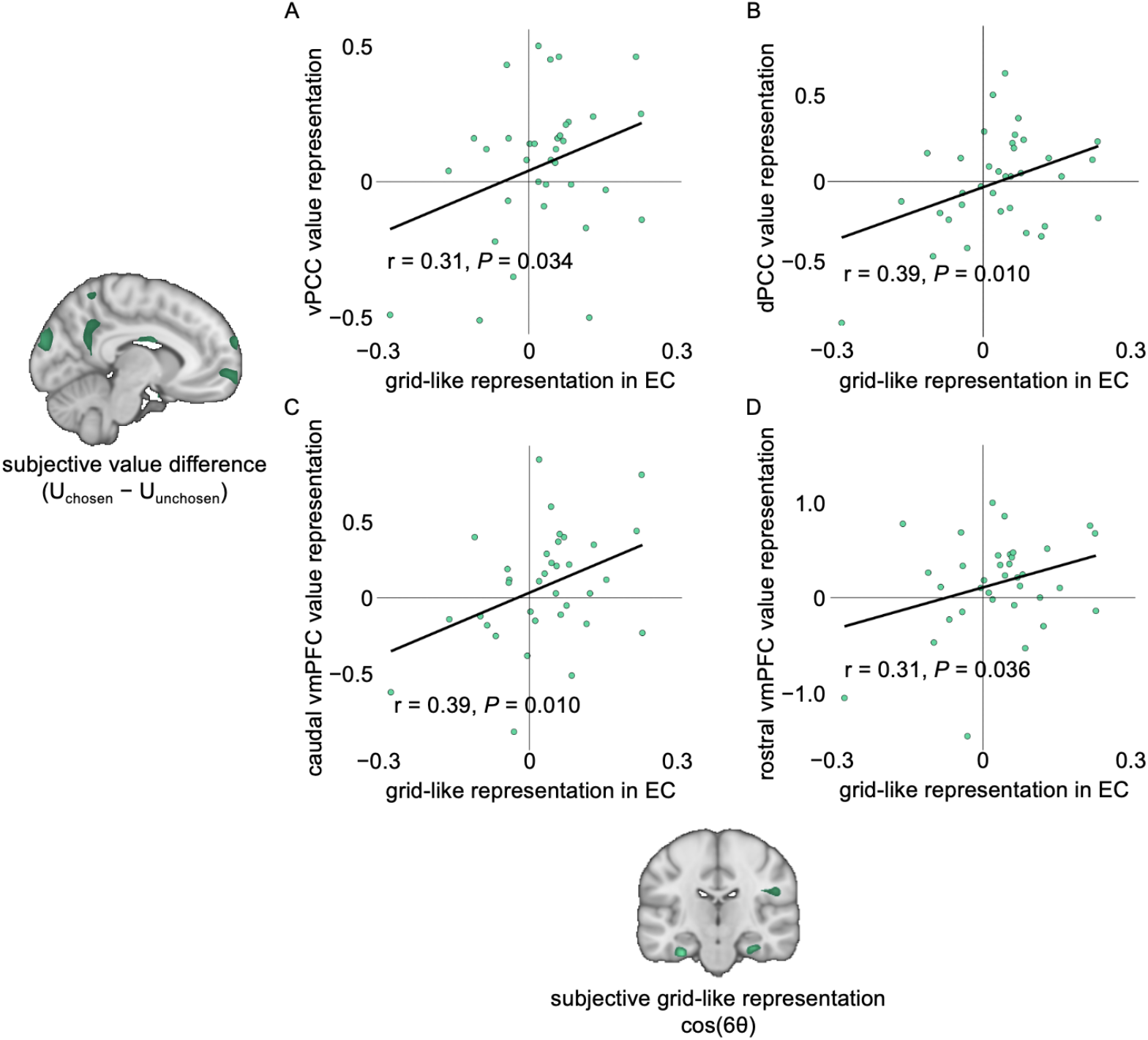
Subjective grid-like and subjective value representations are used in tandem. **A– D)** Across the independently defined value ROIs in the subjective GLM_value_ (i.e., representations of the chosen (relative to unchosen) option) (Fig. 1D, bottom), we observe significant correlations with the subjective grid-like representations (cos(6θ); Fig. 3A, right) in independently defined right EC: vPCC (A), dPCC (B), caudal vmPFC (C), and rostral vmPFC (D).

To examine if grid-like representations are related to individuals’ behavior, we tested if individual estimates of subjective grid-like coding are related to the relationship between Euclidean distance and RT (distance behavioral effect). If so, we expected that the more of an influence Euclidean distance had on RT, the stronger the grid-like representation should be. As hypothesized, we observed a significant correlation between the distance behavioral effect and the subjective grid-like representation across participants in right EC (r = 0.33, *P* < 0.05, one-sided Pearson’s r, Fig. 3D) and mPFC (r = 0.49, P < 0.05, one-sided Pearson’s r). This correlation suggests a strong link between the neural representation of decision vectors over cognitive maps and their use in the decision-making process.

### The EC and mPFC form an abstracted 2D “positional code” of subjective value space

If indeed the EC and mPFC form a cognitive map of an abstract value space, we reasoned that we should observe a positional code that generalizes over stimulus sets ^12,14^. We reasoned that for such a 2D positional code, neural representations for shapes closer together in the value space should be progressively more similar than shapes further away in the abstract space (Fig. 4A). To test our hypothesis, we use representational similarity analysis (RSA) and specified a representational dissimilarity matrix (RDM) that computed the Euclidean distances between shape options defined over a 2D attribute space to explain variance in pattern dissimilarity of BOLD responses to the shape stimuli. To ensure sufficient trial sampling, we combine the shapes into groups of four based on their combined attributes in the value space (Fig. 4A and Extended Data Fig. 7). Further, to ensure we capture abstract representations of the value space that are not confounded by stimulus characteristics within a shape set, we compare activity patterns across shape sets. In other words, we compare neural patterns during viewing of the ovals against neural patterns during viewing of the triangles.

We performed a whole-brain searchlight analysis testing the hypothesized relationship for both objective and subjective value spaces. For both the objective and subjective value spaces, we observed effects of Euclidean distance between options’ positions across shape sets in independently, anatomically defined EC and mPFC: left EC (objective: *P*_TFCE, SVC_ = 0.069, [−34, 2, −36]; subjective: *P*_TFCE, SVC_ = 0.062, [−34, 2, −36]), right EC (objective: *P*_TFCE, SVC_ < 0.05, [26, 6, −42]; subjective: *P*_TFCE, SVC_ < 0.05, [26, 6, −40]), and mPFC (objective: *P*_TFCE, SVC_ < 0.05, [−6, 36, −10]; subjective: *P*_TFCE, SVC_ < 0.05, [14, 38, 14]) (Fig. 4B). Thus, a key component of a cognitive map—an abstract 2D positional code that generalizes over stimulus content—is utilized to represent a value space in EC and mPFC.

We next sought to test whether the subjective 2D positional code is more strongly represented than the objective 2D positional code, as we found with the grid-like representation. First, we compared the 2D positional codes for the objective and subjective RSAs (top 33% of voxels for each individual for each map). We found that in left (*P* < 0.05, one-sided t-test) and right EC (*P* < 0.05, one-sided t-test), and mPFC (*P* < 0.05, one-sided t-test), there was stronger representation of the subjective than the objective positional code (Fig. 4C).

We further reasoned that if the 2D positional code indeed most strongly represents the subjective value space, individuals whose value spaces are more distorted should have stronger representations of a subjective than objective positional code. To test this, we separated subjects into ‘low’ and ‘high’ distortion groups based on amount sensitivity and probability weighting parameter estimates and measured the difference between RSA parameter estimates in the subjective and objective analyses for the three ROIs (left and right EC and mPFC). As expected, individuals with a more distorted value space have a stronger positional code in the subjective relative to the objective RSA than individuals with less distorted value spaces for left (*P* < 0.05, one-sided t-test; Fig 4D) and right EC (*P* < 0.05, one-sided t-test), and mPFC (*P* < 0.05, one-sided t-test). Together, these results confirm a subjective 2D positional code that reflects individual risk preferences in left and right EC and mPFC.

As with the grid-like representation, we also tested if the 2D positional code was related to the distance behavioral effect. Indeed, for left EC (r = 0.31, *P* < 0.05, one-sided Pearson’s r; Fig 4E), and mPFC (r = 0.48, *P* < 0.05, one-sided Pearson’s r), but not right EC (r = 0.15, *P* = 0.19, one-sided Pearson’s r), these measures were positively correlated. Again, this provides evidence of the behavioral relevance of these cognitive map representations in this task.

### Grid-like and value representations in subjective value are used in tandem in risky decision-making

Given that we observe distinct subjective value coding and subjective grid-like representations in this task, we next ask if and how these representations are related. We compare parameter estimates from subjective GLM_value_ in significant value ROIs and parameter estimates from subjective GLM_grid_ in significant grid ROIs and test if these representations covary across subjects. We hypothesize that if these representations are both recruited in a risky decision-making task, an individual’ s representation of the 2D value space should be positively correlated with the strength of its representation in the value coding system. Across subjects, therefore, we should observe a positive correlation between estimates from these analyses.

Specifically, we tested for a positive correlation between the EC or mPFC grid coding (GLM_grid_) and the vmPFC and PCC value difference (GLM_value_) effects across subjects. We observed a positive correlation between the strength of EC subjective grid coding effect and the value difference effects in PCC (vPCC: r = 0.31, *P* < 0.05; dPCC: r = 0.39, *P* < 0.05; one-sided Pearson’s r; Figs. 5A and 5B) and vmPFC (caudal vmPFC: r = 0.39, *P* < 0.05; rostral vmPFC:; r = 0.31, *P* < 0.05; one-sided Pearson’s r; Figs. 5C and 5D). However, we did not observe significant correlations with the mPFC grid-like representation for PCC (vPCC: r = −0.075, *P* = 0.66; dPCC: r = −0.05, *P* = 0.62; one-sided Pearson’s r) or vmPFC (caudal vmPFC: r = −0.068, *P* = 0.67; rostral vmPFC: r = −0.077, *P* = 0.67; one-sided Pearson’s r). To confirm that the relationship between value coding regions and grid-like representations in EC was not merely related to task engagement or choice consistency, we partialled out the inverse temperature term from the cumulative prospect theory model (β from the softmax function that captures how deterministic individuals’ choices are). Indeed, even after controlling for inverse temperature, the positive associations remain significant in PCC (vPCC: ρ = 0.30, *P* < 0.05; dPCC: ρ = 0.35, *P* < 0.05, partial rho; Figs. 5A and 5B) and vmPFC (caudal vmPFC: ρ = 0.37, *P* < 0.05; rostral vmPFC: ρ = 0.30, *P* < 0.05, partial rho; Figs. 5C and 5D). We also confirmed that this result could not be explained by a correlation between the value and grid regressors by controlling for the subject-level correlation values between the first-level regressors (Extended Data Fig. 5). Collectively, these findings link EC grid coding of decision vectors with vmPFC and PCC value comparison coding, suggesting these neural computations operate in tandem to support value construction and comparison during risky choice.

## Discussion

In this study, we utilized a risky decision-making task to test if individuals spontaneously construct and utilize a cognitive map of an abstract value space comprising dimensions of reward probability and amount. We first showed that this task recruits value coding and decision making regions in the brain including vmPFC, PCC, dmFC, and dlPFC. Next, we showed evidence of a grid-like representation of decision vectors through this value space during choice. Specifically, these grid-like representations were identifiable in both EC and mPFC, and were strongest when accounting for individuals’ subjective value distortions in the abstract value space. Further, an independent RSA analysis identified a 2D positional code of this subjective value space in EC and mPFC, abstracting over the visual features of the stimuli and task context. Both the grid-like representations and positional codes in EC and mPFC were correlated with a behavioral measure of cognitive map utilization, demonstrating their behavioral relevance. Finally, the EC grid-like representations were correlated with value representations in vmPFC and PCC. Together, these results show that the cognitive mapping system is used to represent and likely compute vectors over a 2D subjective value space during risky decision-making.

To our knowledge, this is the first demonstration of a *subjectively transformed* abstract value space cognitive map representation during value-based decisions. Specifically, grid representations of decision vectors and 2D positional coding representations were best explained after accounting for distortions due to individuals’ *subjective* preferences. Further, these representations abstract across shape sets, as shown by the RSA analysis. Taken together, this means that the underlying abstract value space maps being encoded are representing “cognitive” information rather than perceptual information or purely objective Euclidean distance. In other words, individuals’ preferences are reflected in the representation. This suggests a novel potential source from which value coding regions may also represent subjectively distorted values that reflect individuals’ preferences ^1–4^. Our finding that the subjective value difference and grid-like representations are correlated across subjects further suggests the cognitive map representation is used in the decision-making process. These results add to a growing body of work that implicates different brain systems in subjective distortions including numerosity tuning in parietal cortex ^49^, event probability encoding in intraparietal cortex and dlPFC ^50^, and affective processing in insula, thalamus, and striatum ^51–53^. How these different systems may interact to shape cognitive map representations, subjective value coding, and risk preferences represents an exciting future direction. This study adds the cognitive mapping system in EC and mPFC to the growing list of interconnected brain regions that play an important role in value-based decision-making, particularly risky choice.

We identified a behavioral signature of a 2D subjective value space. Namely, as participants made choices with longer decision vectors, they took longer to respond—akin to traversing a physical space where further distances take longer to travel. Importantly, this effect persisted after accounting for subjective value difference and other task-related variables. This is an intriguing finding because on one hand, this is not a physical navigation task, so constraints of physical space including things like traveling speed and distance need not apply. On the other hand, theoretical work on computational properties of grid cells suggest the code is error-correcting, given that errors between samples never exceed a certain threshold ^56^. Thus, even though the traversal is not physical, the same limitations of velocity or (equivalently, because of the theta phase–gated location readout ^55^) Euclidean distance between sample locations may be necessary to preserve the error-correcting nature of the representation. This slowing effect was also correlated with neural measures of cognitive map representations across subjects, where cognitive map representations increased with the slowing effect. First, this gives more credence to the behavioral effect representing a cognitive function. Second, it shows that this Euclidean distance and RT analysis provides a robust behavioral proxy of cognitive map representations that may allow these types of representations to be more easily assessed in future work (and can also be applied to some past behavioral studies where cognitive map representations were thought to be unable to be assessed).

The grid-like representation of a subjective value space was present even when only examining never-before-seen decisions (Extended Data Fig. 6). This represents a key theoretical advantage of using a cognitive map representational scheme—it can allow for rapid generalization and zero-shot learning ^8,9,57^. In the present task, choices were made from a large set of novel shapes and required mapping of stimulus dimensions onto the decision-relevant information (reward amount and probability). These features are reflective of many real-world decisions where items are often new and their attributes must be inferred (e.g., a berry’s ripeness based on its color), and may favor the recruitment of the cognitive mapping system.

Our findings show that across subjects the subjective value abstract space representation in EC is correlated with the subjective value difference representation in value coding regions such as vmPFC. While this suggests that these systems work together in subjective valuation, the interplay between these systems in the context of risky choice remains an open question. Intriguingly, in one study vmPFC value-coding neurons were shown to depend on the phase of the recorded theta oscillation, such that value coding occurred when phase-locked to theta and subsequent to grid-coding, suggesting a potential mechanism for interactions between these computations in the same task ^24^. Another important piece of evidence comes from a study showing that causally disrupting hippocampal theta band activity in a value-based decision-making task in non-human primates disrupts reward-based learning and the organization of value coding in OFC ^58^. This and other work provides a key insight that the organization of representations in mPFC and OFC can be driven by theta-locked medial temporal lobe inputs during learning, memory, and decision making ^59–61^. Taken together with our findings here, one way in which subjective value could be constructed from a cognitive map is via EC–HC to mPFC theta band synchrony. Further work using techniques with greater temporal precision is necessary to answer this question in humans.

This work builds on recent findings of grid-like representations of value spaces in non-human primates ^23,24^. First (and perhaps most obviously), it shows the phenomenon of cognitive map formation in value space extends to humans. Second, while Bongioanni and colleagues ^23^ did not find a significant grid-like representation in EC using fMRI and Veselic ^24^ did not record electrophysiological signal from EC, we did observe grid-like representation in EC. Bongioanni and colleagues ^23^ note that the signal-to-noise ratio was weaker in EC than regions which did have grid-like representation of value space (i.e., vmPFC), suggesting that these seemingly disparate results could just be limitations of the available data. Third, one interesting difference that was tested across species is that in both non-human primate studies the grid-like representations showed that the objectively defined, rather than subjectively defined, value space better explained the neural data ^23,24^. While this could represent a species difference, the nature of the stimuli used or the type and amount of training necessary to achieve adequate accuracy in non-human primate tasks could also explain these differences. Future work in both human and non-human primates is needed to better understand what neural components are distinct in humans vs shared across species in cognitive maps and value-based decision-making.

One study examining previously acquired data in an intertemporal valuation task between a default smaller sooner reward and a variable larger later reward, tested for, but did not observe, grid-like representations ^22^ (though, note that in the manuscript they did not report testing for a grid-like representation of a *subjectively* transformed space). One possible explanation for these seemingly disparate results is that the implementation of the cognitive mapping system may be dependent on decision context. For example, situations such as ones with many novel decisions and a large shape set (like our task, as discussed further above) may recruit this system more heavily than situations where an option remains on-screen and can be calculated and compared to a default option. Alternatively, the use of numbers may favor recruitment of the numerosity system, rather than the cognitive mapping system. However, there have not been any other reports in humans (to our knowledge) either confirming or refuting grid-like representations in other value-based decision-making tasks where decisions are made based on subjective preference. Further work is needed to understand the precise circumstances in which the cognitive mapping system is (or is not) utilized in value-based decision-making.

Many of our decisions rely on subjective valuation of different options. Understanding the mechanisms by which individuals form preferences and compute values from decision attributes is therefore crucial for understanding decision-making. This study reveals the previously unknown role of the cognitive mapping system in the subjective valuation process, exposing a new avenue for understanding how value information is constructed and utilized across individuals in decision-making.

## Methods

### Participants and general study procedures

We recruited a total of 98 University of California, Davis (UCD) undergraduate students through the paid SONA recruitment system to participate in this study (age = 22.4 ± 2.52, 72 female, 23 male, 3 non-binary) who did not report having any neurological or psychological conditions. All participants underwent informed consent and the study was approved by the UCD IRB. This study was split into four parts: in-person instruction and training, online training, and two fMRI sessions on separate days. The data from the second fMRI session is not analyzed in this paper. Participants were required to reach a performance threshold in order to participate in the fMRI sessions (***Training procedures***). A high performance threshold was used to ensure that if individuals were representing the shapes using a cognitive map, it would be able to be accurately measured. Forty-one of the 98 participants reached the threshold to complete the fMRI sessions. An additional three participants dropped out before the fMRI sessions, two participants were excluded due to excessive movement in the scanner (assessed during the scanning session), and one participant was excluded due to scanner artifact. Thus, 35 participants (age = 22.4 ± 2.29; 23 female, 10 male, and 2 non-binary) were included in the final analyses.

### Shape ‘sets’

The main fMRI task relied on participants to know the precise probabilities and amounts associated with two sets of shapes. The two sets of shapes were ovals and triangles, respectively. In each set, the shapes had two distinct properties that continuously varied along with probability and amount (Fig. 1B). Specifically, the ovals varied in their width/height ratio and the frequency of the checkerboard pattern inside. The triangles varied in the width of the base and the direction they point. There were three elements of shape sets that were counterbalanced across participants: 1) whether they were first trained on ovals or triangles; 2) whether they were trained on amount or probability first; 3) whether width and mesh or pointing direction and length/width ratio corresponded to probability, leading to 8 total (2×2×2) possibilities. Based on the probabilities and amount steps, there were 100 of each shape per set (10 probability steps × 10 amount steps). However, to reduce the number of choices which include dominated or dominating shapes we never showed shapes from the upper or lower corners of the space (Fig. 2A), thus there were 88 possible shapes from each space that were shown. More specifically, we hypothesized that decisions about these shapes could more easily rely on heuristics (e.g., always choose shapes with high probability and amount). Also, note that participants never saw the space as it is shown in Fig. 2A and instead were taught about each dimension piecemeal and only ever saw one or two shapes at a time during training.

### Training procedures

The goal of the training procedure was for participants to precisely learn the two value dimensions (probability and amount) associated with two sets of shapes (ovals and triangles). To introduce participants to these dimensions, we had them play four training games. Throughout these games, their goal was to earn as much money as possible to get as high as they could on a leaderboard (though, in the training games the money was fictive, and participants were not paid based on performance). Participants played all four games for one shape set before moving on to play the games again for the other shape set (the order for whether ovals or triangles were learned first was counterbalanced).

#### Game 1

In game one, participants were allocated 10 tokens. They were told they would be shown 20 shapes and asked to choose whether or not to use a token. The tokens multiplied the amount of each shape by 10. At the end of each choice, a uniform random number was drawn between 0 and 1, and if the probability was lower than the number, participants won the amount associated with the shape (×10 if they used a token). Periodically, they were asked questions about what dimension corresponded to the stimulus characteristics (e.g., to which does the direction that the triangle is pointing correspond: probability or amount?). The goal of this game was to get participants familiar with the shapes and start to associate the stimulus characteristics with probability and amount.

#### Game 2

In game two, participants were shown two shapes and asked to choose which shape was greater in a given dimension (either probability or amount). Each shape was only different by one ‘step’ (i.e., 10% or $1). They were told after each choice whether or not they were correct. This game was not paid. The goal of this game was to help participants learn to discriminate between stimuli at the smallest difference they would see.

#### Game 3

In game three, participants were asked to select (from $1–10 or 10–100%) the amount or probability associated with a given shape. After each choice they were given feedback about the true answer and how close they were. They were not paid based on performance in this game. The goal of this game was to ensure participants were accurate in knowing the precise amounts and probabilities of each shape. Unbeknownst to participants, performance in this game was used to evaluate whether participants qualified for the fMRI session (see below for further details).

#### Game 4

In game four, participants were shown two shapes and asked to choose one of the shapes. After choosing a shape, the probabilities and amounts associated with both shapes were shown. Then, the probabilities were played out, and, if successful, participants were awarded the money associated with the chosen shape. The purpose of this game was to have participants begin to think about comparing shapes.

We additionally had them play a fifth game which was a more classical neuroeconomic–style risky decision-making task (not involving the shapes) where probabilities and amounts were explicitly shown (in pie form, and text, respectively) and their amount sensitivity parameter estimates were acquired using a staircase procedure. This was initially included to get an additional measure of amount sensitivity but is not analyzed in this paper.

At the end of each training session (i.e., all five games), participants were shown text indicating whether they qualified for the fMRI task for that shape set. Finally, participants were shown their place on a leaderboard with the amount of money they acquired throughout the games compared to other participants. This leaderboard was initialized with placeholder values but was updated after each in-person training session with each real score attached to an anonymized name. In order to attend the fMRI sessions, participants were required to qualify for both shape sets. They were not aware of the specific qualification criteria, but were aware that it was based on how well they knew the probabilities and amounts associated with each shape dimension. In actuality, to qualify participants needed to get within an average of 0.5 ‘steps’ in game three (i.e., less than 5% or $0.50). Throughout all the training games, for each set, a selection of 10 shapes was selected to show the full range of possibilities for a given dimension (which included one shape which repeated for both dimensions; Fig. 1B) and 12 other shapes. Thus, only 31/88 (19+12, 35%) shapes per set were ever viewed before the fMRI task (Extended Data Fig. 8).

### In-person training

The first part of the experiment was in-person training. At this visit, participants were consented, given information about the structure of the whole study, and then instructed-on and began the in-person training session. This involved completing the training session for each shape set once. Participants were paid $10 for completion of this in-person training session.

### Online training

After the in-person training, if participants did not reach the required performance level on both shape sets they were asked to continue training online (twelve participants qualified during the first in-person session). They completed the online training at their own pace from a web browser and were paid $2.50 for each completed session up to five sessions (per shape). Participants were scheduled for the fMRI session after meeting qualification criteria for both sets, but could still get paid for completing additional training sets (up to five).

### Shape preference task

After meeting qualification criteria (***Training procedures***), participants were invited to the lab to complete the shape preference task while undergoing fMRI scanning. Participants completed the training session once more the day before or day of scanning. The shape preference task included three runs. In each run, participants were asked to make a series of binary choices between shapes. Shapes were shown sequentially to participants (Fig. 1B) and they were asked to choose which shape they preferred, the ‘previous’ or ‘current’ shape. Each choice was between two shapes from the same set. Participants were given 5s for each choice; if they took longer the choice was recycled and presented again at the end of the run. Each run included 70 choices, plus six additional practice trials identical to trials in training game four for the first run, thus including 216 choices total (6 + 70 choices x 3 runs). However, the practice trials were not included in the fMRI analysis. The choices were incentive compatible such that at the end of each run, one of the chosen shapes was selected, a number was uniformly drawn from 0-1, and if the number was less than the probability associated with the shape, participants were given the amount associated with that shape.

Participants were paid $30 for each scanning session, plus this performance-based bonus of $0– 30.

### fMRI scanning procedure

We used a Siemens Skyra 3 Tesla scanner. Our T2* functional images were acquired with a multiband (acceleration factor = 2) echo-planar-imaging (EPI) sequence, at a 30º angle with the AC-PC line ^62^. The parameters used were as follows: 46 slices, 3 mm thick, repetition time (TR) = 1300 ms, echo time (TE) = 24 ms, flip angle (FA) = 67º, field of view (FoV) = 192 mm, voxel size = 3 mm isotropic. We acquired T1-weighted images to use in the normalization process. Our T1-weighted images were acquired using a magnetization-prepared rapid gradient echo (MPRAGE) sequence: TR = 1800 ms, TE = 2.96 ms, FA = 7º, FoV = 256 mm, voxel size = 1 mm isotropic. To account for field inhomogeneities we acquired images using a gradient recalled echo (GRE) sequence: 42 slices, TR = 630 ms, TE_1_ = 10 ms, TE_2_ = 12.46 ms, FA = 40º, FoV = 192 mm, voxel size = 3 mm isotropic.

### fMRI preprocessing

Results included in this manuscript come from preprocessing performed using fMRIPrep 22.0.1 (^63^; RRID:SCR_016216), which is based on Nipype 1.8.4 (^64^; RRID:SCR_002502).

#### Preprocessing of B0 inhomogeneity mappings

A total of 2 fieldmaps were found available within the input BIDS structure for this particular subject. A B0 nonuniformity map (or fieldmap) was estimated from the phase-drift map(s) measure with two consecutive GRE (gradient-recalled echo) acquisitions. The corresponding phase-map(s) were phase-unwrapped with prelude (FSL 6.0.5.1:57b01774).

#### Anatomical data preprocessing

A total of 2 T1-weighted (T1w) images were found within the input BIDS dataset. All of them were corrected for intensity non-uniformity (INU) with N4BiasFieldCorrection ^65^, distributed with ANTs 2.3.3 (^66^, RRID:SCR_004757). The T1w-reference was then skull-stripped with a Nipype implementation of the antsBrainExtraction.sh workflow (from ANTs), using OASIS30ANTs as target template. Brain tissue segmentation of cerebrospinal fluid (CSF), white-matter (WM) and gray-matter (GM) was performed on the brain-extracted T1w using fast (FSL 6.0.5.1:57b01774, RRID:SCR_002823, ^67^). A T1w-reference map was computed after registration of 2 T1w images (after INU-correction) using mri_robust_template (FreeSurfer 7.2.0, ^68^). Brain surfaces were reconstructed using recon-all (FreeSurfer 7.2.0, RRID:SCR_001847,), and the brain mask estimated previously was refined with a custom variation of the method to reconcile ANTs-derived and FreeSurfer-derived segmentations of the cortical gray-matter of Mindboggle (RRID:SCR_002438, ^69^). Volume-based spatial normalization to one standard space (MNI152NLin2009cAsym) was performed through nonlinear registration with antsRegistration (ANTs 2.3.3), using brain-extracted versions of both T1w reference and the T1w template. The following template was selected for spatial normalization: ICBM 152 Nonlinear Asymmetrical template version 2009c [^70^, RRID:SCR_008796; TemplateFlow ID: MNI152NLin2009cAsym].

#### Functional data preprocessing

For each of the 6 BOLD runs found per subject (across all tasks and sessions), the following preprocessing was performed. First, a reference volume and its skull-stripped version were generated using a custom methodology of fMRIPrep. Head-motion parameters with respect to the BOLD reference (transformation matrices, and six corresponding rotation and translation parameters) are estimated before any spatiotemporal filtering using mcflirt (FSL 6.0.5.1:57b01774, ^71^). The estimated fieldmap was then aligned with rigid-registration to the target EPI (echo-planar imaging) reference run. The field coefficients were mapped on to the reference EPI using the transform. BOLD runs were slice-time corrected to 0.612s (0.5 of slice acquisition range 0s-1.23s) using 3dTshift from AFNI (^72^, RRID:SCR_005927). The BOLD reference was then co-registered to the T1w reference using bbregister (FreeSurfer) which implements boundary-based registration ^73^. Co-registration was configured with six degrees of freedom. Several confounding time-series were calculated based on the preprocessed BOLD: framewise displacement (FD), DVARS and three region-wise global signals. FD was computed using two formulations following Power (absolute sum of relative motions, ^74^) and Jenkinson (relative root mean square displacement between affines, ^71^). FD and DVARS are calculated for each functional run, both using their implementations in Nipype (following the definitions by ^74^). The three global signals are extracted within the CSF, the WM, and the whole-brain masks. Additionally, a set of physiological regressors were extracted to allow for component-based noise correction (CompCor, ^75^). Principal components are estimated after high-pass filtering the preprocessed BOLD time-series (using a discrete cosine filter with 128s cut-off) for the two CompCor variants: temporal (tCompCor) and anatomical (aCompCor). tCompCor components are then calculated from the top 2% variable voxels within the brain mask. For aCompCor, three probabilistic masks (CSF, WM and combined CSF+WM) are generated in anatomical space. The implementation differs from that of Behzadi et al. in that instead of eroding the masks by 2 pixels on BOLD space, a mask of pixels that likely contain a volume fraction of GM is subtracted from the aCompCor masks. This mask is obtained by dilating a GM mask extracted from the FreeSurfer’s aseg segmentation, and it ensures components are not extracted from voxels containing a minimal fraction of GM. Finally, these masks are resampled into BOLD space and binarized by thresholding at 0.99 (as in the original implementation). Components are also calculated separately within the WM and CSF masks. For each CompCor decomposition, the k components with the largest singular values are retained, such that the retained components’ time series are sufficient to explain 50 percent of variance across the nuisance mask (CSF, WM, combined, or temporal). The remaining components are dropped from consideration. The head-motion estimates calculated in the correction step were also placed within the corresponding confounds file. The confound time series derived from head motion estimates and global signals were expanded with the inclusion of temporal derivatives and quadratic terms for each ^76^. Frames that exceeded a threshold of 0.5 mm FD or 1.5 standardized DVARS were annotated as motion outliers. Additional nuisance timeseries are calculated by means of principal components analysis of the signal found within a thin band (crown) of voxels around the edge of the brain, as proposed by ^77^. The BOLD time-series were resampled into standard space, generating a preprocessed BOLD run in MNI152NLin2009cAsym space. First, a reference volume and its skull-stripped version were generated using a custom methodology of fMRIPrep. All resamplings can be performed with a single interpolation step by composing all the pertinent transformations (i.e. head-motion transform matrices, susceptibility distortion correction when available, and co-registrations to anatomical and output spaces). Gridded (volumetric) resamplings were performed using antsApplyTransforms (ANTs), configured with Lanczos interpolation to minimize the smoothing effects of other kernels ^78^. Non-gridded (surface) resamplings were performed using mri_vol2surf (FreeSurfer).

Many internal operations of fMRIPrep use Nilearn 0.9.1 ^79^, RRID:SCR_001362), mostly within the functional processing workflow. For more details of the pipeline, see the section corresponding to workflows in fMRIPrep’s documentation.

#### Additional Preprocessing

Subsequent to the fMRIprep preprocessing, we smoothed BOLD images using an 8mm full width at half maximum (FWHM) kernel. These images were used for all univariate analyses while unsmoothed images were used for RSA.

### Response time regression analyses

To understand the role of Euclidean distance on RT, we performed a two-step mixed effects regression. Specifically, for step one we z-scored the natural log of the response times and |subjective value difference| separately for each participant and then estimated a mixed effects regression of |subjective value difference| on RT, specifying participant as a random effect. At step two, we regressed the z-scored (within participant) Euclidean distance between shapes on the residuals from step one. For analyses examining correlations between neural representations (grid-like and positional code) and the behavioral distance effect, we used the same two-step procedure but estimated separate regressions for each participant (and thus, did not specify any random effects for either step).

### Computational Modeling

We utilize two computational models in this task, the objective value model and the subjective value model. These models are utilized to predict individuals’ shape choices in the shape preference task. The objective value model is a nested version of the subjective value model (with two of the three parameters fixed). We first describe the subjective value model based on cumulative prospect theory ^34^.

#### Subjective value model

We first estimate the subjective value of each shape using nonlinear transformations for amount and probability:

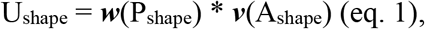

where P and A represent the probability and amount associated with the shape, ***w***(P) represents the probability weighting function ^80^:

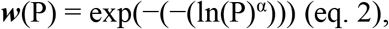

Where 0 < α < 1 reflects overweighting low probabilities and underweighting high probabilities, and vice versa for α > 1 and ***v***(A) represents the power utility function ^81^:

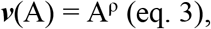

where 0 < ρ < 1 reflects risk aversion and ρ > 1 reflects risk seeking. The difference between shape utilities was calculated,

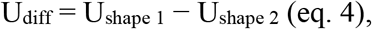

then estimated subjective values for each shape were then passed through a softmax function:

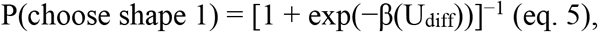

where β is an inverse temperature parameter that reflects decision consistency.

#### Objective value model

The objective value model takes the same form as the subjective value model except α and ρ are fixed to 1. Effectively, this means that the value of a given shape is simply

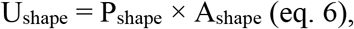

where there is no transformation on probability or amount. Thus, the only free parameter in this model is the inverse temperature parameter.

### Computational Model Specification

For both models, 216 trials per participant were used to estimate parameters. We utilized hierarchical Bayesian estimation ^82^ where all parameters for each subject were drawn from a group level distribution. Wide priors were specified for the group-level mean and standard deviation parameters: Normal(0,10) and Cauchy(0,2.5), respectively. For efficient sampling, we used non-centered parameterizations ^83^ (sometimes referred to as the ‘Matt trick’) where individual parameters are specified as coming from a Normal(0,1) distribution, and multiplied by the group standard deviation and added to the group mean. Subsequently, these values were passed through the Phi_approx function (a computationally efficient approximation of the cumulative normal distribution) and multiplied by 50, 2, and 10, for the inverse temperature (β), power utility (ρ), and probability weighting (α) parameters, respectively, constraining them between 0 and these values.

### Computational Model Comparison

To compare between the two models, we used the integrated Bayesian Information Criterion (iBIC; ^36^). The iBIC allows for comparing models where an entire posterior distribution is estimated rather than just point estimates. Because these models are hierarchical, each estimated parameter effectively has two freely estimated parameters (mean and variance). Thus, the iBIC ‘penalizes’ the subjective model for having 6 parameters (2 × 3 [β, ρ, and α]) and the subjective value model for 2 parameters (2 × 1 [β]).

### Computational Model fitting procedure

To fit the model parameters we used the R implementation ^84^ of Stan ^85^. We used the default sampling procedure, the No-U-Turn Sampler (NUTS) ^86^ which is a variant of the Hamiltonian Monte Carlo (HMC) estimation procedure. For each model, we ran four chains which estimated 5000 samples each. We specified the first 2000 samples for burn-in, and after discarding these obtained 12,000 usable samples. All values of 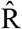, the potential scale reduction factor, were below 1.1, all effective Ns were > 2600, and the chains were all visually inspected to ensure good mixing and convergence ^87^. For each individual and each model, the median estimated parameters were utilized.

### Measurement of subjective value space distortion

To measure the extent to which participants’ subjective value spaces were distorted, we examined estimated computational model parameters. First, we measured the distance from the non-distorted objective value space:

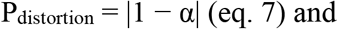

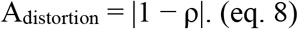

Because parameter values of 1 represent the non-distorted objective dimensions, the greater deviance from one, the more the space becomes distorted in either dimension. After obtaining the distortion separately for each dimension, we separately normalize the values for probability and amount distortion each such that 0 represents the least and 1 represents the most distorted subject. Finally, we averaged the normalized P_distortion_ and A_distortion_ values for each subject to obtain a measure of overall distortion based on both dimensions. This value is used to group participants for the RSA analysis testing representation strength for the objective relative to subjective positional code (i.e., Fig. 4C).

### fMRI Analysis

#### Univariate General Linear Models (GLMs)

We utilized two GLMs to test for a value difference signal (GLM_value_) and grid-like representation of decision vectors (GLM_grid_). For both of these analyses, we analyzed BOLD signal at the time of the second shape appearing. Both GLMs were set up in the same way, with the exception of the parametric modulator at the time of the second shape. First, we will describe the commonalities in the GLMs and then specify how the parametric modulator differed between GLM_value_ and GLM_grid_. The regressors included the following, convolved with the hemodynamic response function (HRF):

1. *Shape 1 onset (duration 2s)*: the onset of the first shape,
2. *Shape 2 onset (0s)*: the onset of the second shape,
3. *Shape 2 onset parametric modulator (0s):* either the value difference or grid-like modulation regressor (see below for further details) at the time of onset of the second shape,
4. *Choice, shape 1 (0s)*: the onset of a choice, only when shape 1 was chosen,
5. *Choice, shape 2 (0s)*: the onset of a choice, only when shape 2 was chosen.

For GLM_grid_ we specified these regressors separately for shape set 1 and shape set 2, resulting in 10 total regressors. Additionally, we added regressors (not convolved with the HRF) to censor volumes with TR-by-TR framewise displacement (FD) > 0.9 mm (one regressor for each censored volume; ^88^).

For GLM_value_, the parametric modulator on shape 2 onset was specified as U_chosen shape_ − U_unchosen shape_. We estimated an objective and subjective GLM_value_, where the objective and subjective models were used to calculate U, respectively.

For GLM_grid_, the parametric modulator on shape 2 onset was cos(6θ), where θ is the angle of the decision vector through the value space. As with GLM_value_, we estimated an objective and subjective GLM2 using the angles through the objective and subjective value spaces, respectively. Multiplying by angle by six before taking the cosine specifies the canonical six-fold symmetry exhibited by grid cells activity. In control analyses (Fig. 3C) we estimated the GLM using angles in subjective space, but instead using 4, 5, 7, and 8 to specify four-, five-, seven-, and eight-fold symmetries, respectively. Note that some prior work uses left out data to estimate a ‘preferred’ angle to which the grid-like representation is oriented. However, in this work we do not implement this procedure given animal ^26,29^ and recent human work ^89–92^ showing that ‘preferred’ grid orientations anchor to relevant behavioral dimensions. Here, angles are calculated with respect to the probability dimension.

SPM12 was used to estimate the models and a one-sample t-test was used to compare across subjects at the second level. All parametric modulators were z-scored for each participant. We estimated all three runs together with a separate intercept for the mean difference in each run.

We performed whole-brain and small volume–corrected threshold-free cluster-enhancement (TFCE) ^93^ with 1000 permutations to determine significance. Additionally, to further analyze the grid-like effects for each ROI one-sided paired-sample t-tests were used to compare between groups (e.g., objective vs subjective grid-like representation). The values extracted from the ROIs were the mean values across participants.

We selected two sets of ROIs for the univariate fMRI analyses: value coding and grid-like. The value coding ROIs were defined as 10 mm radius spheres from two peaks in vmPFC and two peaks in PCC taken from a meta analysis examining regions of the brain responsive to value (Figure 1 and Table 3 from ^2^): caudal vmPFC, rostral vmPFC, dorsal PCC (dPCC), and ventral PCC (vPCC). The grid-like ROIs were taken from a previous study identifying grid-like representation in an abstract navigation task ^12^. In that study, these areas were identified using an F-test procedure and thresholded within anatomical ROIs and at *P* < 0.01. This resulted in a right EC ROI and an mPFC ROI. Additionally, to look at left EC, we multiplied the x location of the right EC ROI by −1.

To confirm that our results were specific to the six-fold symmetry inherent in grid cells, we tested different versions of the subjective GLM2 using the regressors cos(4θ), cos(5θ), cos(7θ) and cos(8θ), to specify four-, five-, seven-, and eight-fold symmetries, respectively. We then extracted the data from each ROI, for each participant, after further thresholding to average across only the voxels with *P*_uncorrected_ < 0.05 in the group-level grid-like analysis. A one-sided t-test (> 0) was then performed for each symmetry.

#### Representational Similarity Analysis (RSA)

We utilized RSA ^94^ to assess patterns in BOLD response to shapes in our task. In the RSA model, we sought to examine if patterns for shapes in more similar locations in the abstract value space were more similar than shapes in more distant locations in the abstract value space. Because there were 88 shapes that could be viewed for each shape set, we grouped shapes into sets of four. For example, shapes with [$5 or $6] and [30% or 40%] were placed into one group to increase examples in each category while also being relatively specific to the location in abstract space (Extended Data Fig. 7). This resulted in 23 groups per shape set. We then estimated GLMs with separate regressors for each of these groups, for each shape set. Each regressor included the onset of the shape (whether it was first or second presented) for a 2s duration. The GLM also included separate regressors for *choice, shape 1* and *choice, shape 2* and censored volumes with FD > 0.9 mm (as with the univariate analyses). A t-contrast was performed for each shape group. Note that this GLM was performed on unsmoothed data.

We performed two analyses— one for the objective value space and one for the subjective value space. These were identical except for how the Euclidean distance between stimuli sets were calculated. For every voxel in the whole-brain mask, we extracted t-statistics from a 100 voxel– sized sphere. These t-statistics were then vectorized and z-scored. These z-scored values across all groups for shape set 1 were then compared to all the z-score values across all groups in shape set 2 using Euclidean distance. By comparing across shape sets we are able to 1) compare across visually distinct stimuli, such that the only shared characteristic between them is the location in an abstract map, and 2) Compare across trials to avoid any temporal autocorrelation that could result in inflated results. The resulting 23 × 23 neural RDM was formed by subtracting these values from 1 (to convert similarity to dissimilarity). Finally, the upper triangle of this RDM (including the diagonal) was obtained by averaging the bottom and top triangles.

The model RDM was created by taking the mean of each group (i.e., the [$5 or $6], [30% or 40%] group was $5.50 and 35%). In the case of the subjective RSA, this mean was distorted according to each individuals’ amount sensitivity and probability weighting parameters. The 2D Euclidean distance was then calculated between all other group means. The upper triangle of this RDM was compared to the upper triangle of the neural RDM using Pearson’s r to get a similarity score. Note that Pearson’s r was used to enable the comparison between subjective and objective. A rank-based measure (such as Kendall’s tau) would not be able to differentiate between these two RDMs. To ensure this did not cause bias in our results, we used non-parametric permutation testing to assess significance. Specifically, we additionally shuffled the values of the model RDM in 1000 permutations to construct the null distribution. The average value across the permutations was subtracted from the similarity score for each searchlight voxel. The resulting map was then smoothed at 8mm FWHM. To assess significance for the searchlight method, after obtaining a value for each voxel, a t-test was performed across all subjects. TFCE was used to account for multiple-comparisons.

Subsequent to the whole-brain analysis, we performed several analyses on extracted voxels per subject. To do so, we utilized the top 33% of voxels for each subject for each RSA (subjective and objective) within the specified ROI (as detailed below). To compare the subjective and objective RSA results for each ROI, we performed a paired t-test between the averaged values for the subjective and objective RSA (one sided, for subjective > objective). We further compared the subjective and objective RSA results between the high and low distortion groups, using these same averages using an (unpaired) t-test (one sided for high distortion > low distortion).

We utilized anatomically defined ROIs for small volume correction, including right and left EC from the Jülich atlas and mPFC defined by combining right and left 32d and 32pl from the Neubert atlas ^95^.

## Supporting information

Experiment Instructions

## Data and code availability

All the data necessary to reproduce the findings of this study are available on Open Science Framework at osf.io/yxm7s. The unthresholded group-level statistical parametric maps are available on NeuroVault at identifiers.org/neurovault.collection:21751. Custom code is available upon reasonable request to the corresponding author.

## Author contributions

M.A.O., S.A.P. and E.D.B. conceived the project and designed the experiments. M.A.O. and J.B. recruited and trained participants and collected data. With supervision from E.D.B. and P.D., M.A.O. analyzed data. M.A.O. and E.D.B. wrote the paper. All authors contributed edits and revisions to the paper and approved the final manuscript.

