## Supplementary material for "A cognitive map of subjective value space for human risky choice": Experiment Instructions

### OVERVIEW

The year is 2480. You are a space engineer coming home from a mining expedition. You find yourself waking up from hyper-sleep earlier than expected. The spaceship appears to be making an emergency landing, and you notice that you are landing on an alien planet. As you begin to become more aware of your surroundings and less groggy from the hyper-sleep, you notice an alarm sounding and a robotic voice repeating over the loudspeaker, “error, error, warp drive malfunction, fix immediately.” Your crewmates remain under hyper-sleep per federal regulations. You were selected as the designated emergency responder in case of any malfunctions given your background in spaceship engineering and alien culture studies. You are the only one allowed to be walking around the ship right now, and you are the only one who can save it. You have one and only one goal: Get a warp drive.

You decide to explore the planet in pursuit of a warp drive. As you walk around, you soon notice a billboard advertising an auction. Among the items pictured for sale is a warp drive! You just need to figure out how to get enough money to outbid everyone else to get the warp drive. Soon after seeing the billboard, you come across a building with familiar currency symbols and flashing lights and arrows. You go inside and see human-like creatures standing around two tables. Each table has shapes flashing on a screen. When shapes appear, creatures point at the screen, and are sometimes handed money depending. You realize this is the perfect opportunity to make some money so you can win the warp drive at the auction! *If only you understood how they are choosing which shapes to select…* As you stand near one of the tables trying to understand how the game works, one of the creatures wanders over to you, noticing your confused look, and hands you a sheet of paper with some text (somehow translated to your language) and pictures of shapes matching those in this alien game:

### TRAINING INSTRUCTIONS

There are two main games at this casino. These games are both based on making decisions between shapes. The key to making these decisions is to learn as much as you can about what these shapes mean. We have prepared some instructions and some short training games for you to play to help you learn about these shapes. Please continue reading for these instructions.

Each shape has an ***amount*** (in ***$***) and a ***chance*** (or probability) ***of winning*** that amount. In the example below, the shape shown has a 60% ***chance*** (shown by the green area) of winning ***$***5:

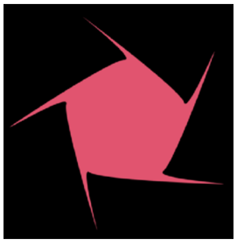
Shape

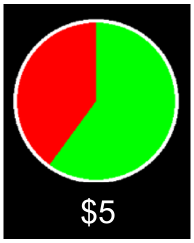

Pie chart

Different aspects of the shape will change according to the ***chances of winning*** and ***amount of $*** of that shape. For example, the color of the shape might change according to the ***amount of $*** of that shape. You will learn what aspects of the shapes change and what ***amounts*** and ***chances*** they represent as you play the training games. There are ten possibilities for the amounts of ***$*** of each shape ranging from $1 to $10 in steps of 1. There are also ten possibilities for the chances of winning for each shape, ranging from 10% to 100%, also in steps of 10.

#### *TRAINING GAME 1*

In this training game, you will see many shapes like that pictured above. This game is further split into two parts. In the first part of the training game, the amount of $ of all the shapes you see will be the *same* ($5), but they will all have *different* ***chances of winning***. In the second part of the training, the chances of winning will be the *same* for all the shapes (60%), but they will have *different* ***amounts of $***. In these training sessions, you will see a 'pie chart' appear after each shape which will indicate that shape's chance of winning and the amount of $ of that shape.

For each shape you see, the gamble will be played out and you will have a chance (proportional to the green area shown in the pie chart) to win the points shown. If you win, you will see the amount appear in **green**. If you lose, a red **0** will appear. When you see each shape, you will have the option to 'use' or 'keep' a token. If you use a token, it will multiply your winnings by 10. You will be given 10 tokens at the beginning of each part. When each part ends, you will lose your unused tokens, so be sure to use them all in each part. You will be completing 20 trials in each part, so you should use your tokens on 50% of the shapes. In other words, you should accept the shapes worth $6 or more and shapes with a 60% or greater **chance of winning**. A counter on the top right corner of the screen will keep track of how many trials you have completed during a given part. An example round where the player used their token and won is shown below:

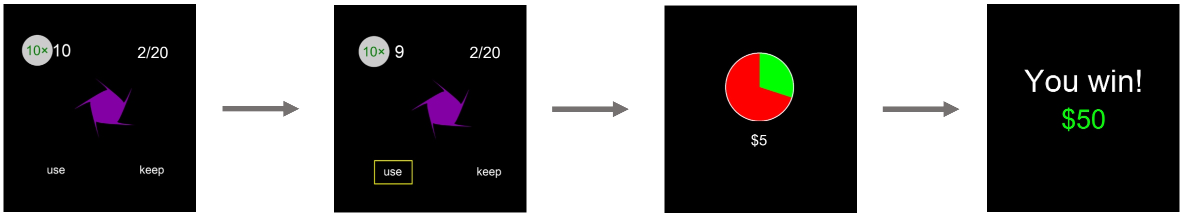


For each part of *training game 1*, you also will periodically get asked questions about the shapes.

*TRAINING GAME 2*

In *training game 2*, you will be asked to make choices between two shapes. These shapes will differ from each other by only one 'step.' For example, in one trial these shapes may have the *same* chance but one shape is worth $6 while the other has is worth $5. In another example trial, the shapes both may have the same amount of $ but different chances of winning, where one has a 30% chance and the other has a 40% chance. An example trial where the player was asked to indicate the shape with the highest probability and answered correctly is below:

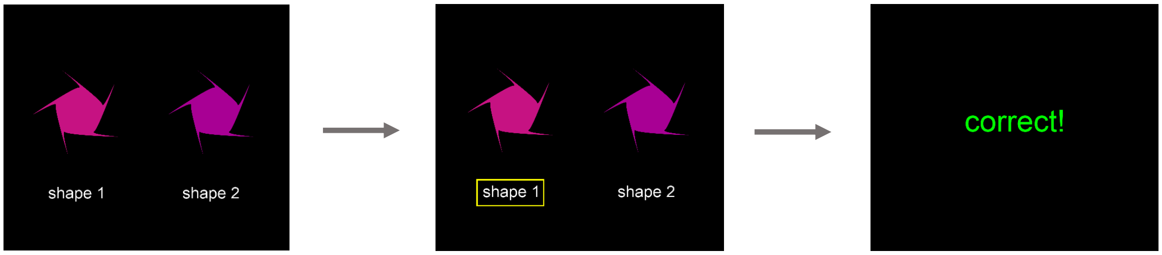


*TRAINING GAME 3*

In *training game 3*, you will be asked to choose the ***chance of winning*** or ***amount of $*** of each shape. You will be shown one shape and asked to choose the correct answer among the options below. If you are wrong, the correct answer will be highlighted in **green** and your choice will be highlighted in **red**. If you are incorrect but on the right track, the correct answer will be highlighted in **green** and your choice will be highlighted in **orange.** If you are close the correct answer will be highlighted in **green** and your choice will be shown in **light green**. If you are exactly right, you will see your choice show up in **green**. An example trial is below where the player was close but not exactly right:

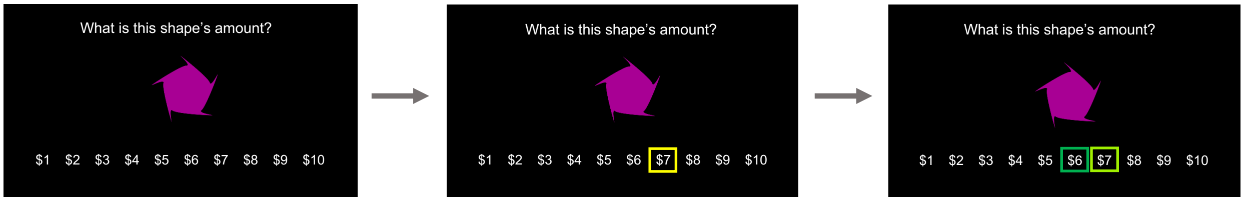


*TRAINING GAME 4*

Next, you will begin *training game 4*. In *training game 4*, you will make a series of choices between two shapes. When making this choice, try to choose the best shape considering both the ***amount***

***of $** and **chance of winning*** for each shape. For each choice, you will win the amount played out from the chosen shape based on its ***amount of $*** and ***chance of winning***. After your choice and before you see your outcome, you will have a chance to see the pie charts underlying each shape. An example round is shown below:

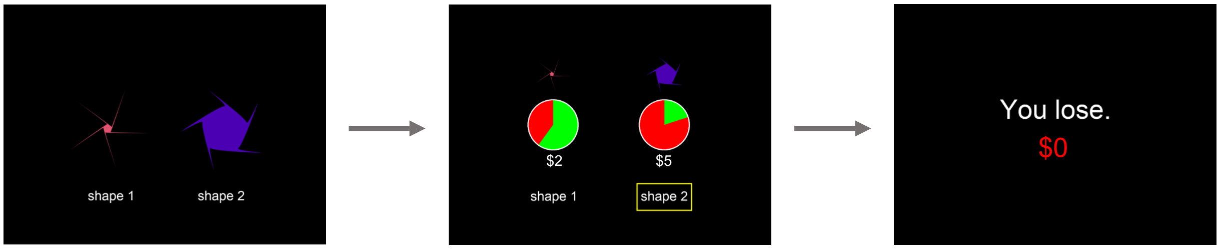


### *TRAINING GAME 5*

In this training game, you will not see the shapes. Instead, you will be asked to make a series of choices between the gambles depicted by the pie charts. You will win the amount played out from the chosen gamble. An example round where someone chose the gamble on the right is shown below:

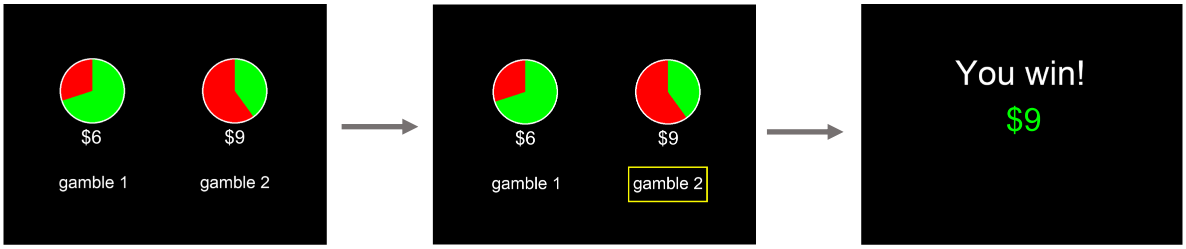


### *FINAL NOTES*

To make choices in these games, you will use two buttons, the ‘A’ and ‘L’ keys on your keyboard, corresponding to ‘left’ and ‘right,’ respectively. Use the ‘left’ button to choose an option on the left and the ‘right’ button to choose an option on the right. For questions where there are more than two answers (i.e., *training game 3*), use the ‘left’ or ‘right’ button to move the box until it is on top of the answer you want (if it begins on top of the choice you want, you will need to move it off and on). After waiting a second it will automatically select that option.

For the first 3 choices in each training game you will have as much time as you need to make a choice. However, the rest of the trials in each game will require you to respond within 6 seconds. If you do not respond within 6 seconds you will see an **X** appear and the choice will be shown to you again at a later point. Please do your best to answer before the time is up.

You will complete all the training games first for one set of shapes, and then will play all of them again with a *different* set of shapes. The two main games at the casino will use both sets of shapes. For each set of shapes, distinct features of the shapes will change depending on the ***chance of winning*** and ***amount of $*** of each shape. It is important to learn these features as best as possible because the two main casino games will require you to simultaneously consider both the ***chance of winning*** and ***amount of $*** of each shape when making decisions. We are keeping track of the high scores from previous participants (with anonymized names), which you will have a chance to see after each session. See how high of a score you can get and how you compare to other participants!

### ONLINE TRAINING INSTRUCTIONS

You have *five* chances to qualify for the MRI portion of this study by completing the online training. This online training consists of the same 5 games you played during your in-person visit. The purpose of the training is to get as familiar as possible with the amount of $ and chance of winning for each shape. Whether or not you qualify for the MRI portion of the study depends on how well you are able to learn the shapes. This will be automatically calculated and revealed to you at the end of the training sessions before you see the leaderboard. Note that you will need to see that you qualify for *both* sets of shapes (i.e., training: set 1 and training: set 2) to participate in the MRI portion. Also please note that learning these shapes to the extent we are requiring is quite difficult, so do not be discouraged if you are unable to learn in the first few sessions or even at all! You will be paid $5 for each time training is completed for both sets (i.e., once training: set 1 *and* training: set 2 are completed, you will earn $5). *Even if you qualify, you can still be paid $5 for each time the training is completed up to 5 times total.* In other words, you will earn $25 if you complete the training sessions five times. Note that this task periodically makes noise, so be sure to adjust your computer's volume appropriately and/or use headphones if necessary. To complete the training, follow these instructions:

1. Go to https://bit.ly/3qI9FhH
2. NOTE: Once the experiment has loaded, your browser will automatically make itself full screen (you can exit full-screen mode by hitting the 'esc' key on your keyboard)
3. Enter your Subject ID when prompted: __________
4. Select 'training: set 1' and click OK to start the task
5. Once you are finished with the task, your browser window will minimize itself
6. Repeat for 'training: set 2'
7. Repeat 1–6 up to five times

If you have any issues or questions, please contact me at.

### GAME 1 OVERVIEW

Now that you have completed the training, you feel confident that you can make choices between the shapes. However, you’re still not exactly sure what to do in the two games to earn money. As if they were reading your mind, another nearby alien creature hands you another sheet of paper in your language. They gesture to an open seat at the table and indicate that it will be waiting for you when you finish reading the instructions:

**GAME 1 INSTRUCTIONS**

In *game 1*, instead of appearing next to each other like in the training, shapes will appear one after the other. After seeing the second shape you will have 5 seconds to decide between the two shapes you were presented with: the previous shape and the current shape. The text 'previous' will refer to the first shape shown and 'current' will refer to the current shape on the screen. You will see shapes from both worlds. An example round is shown below where the player chose the purple 'previous' shape:

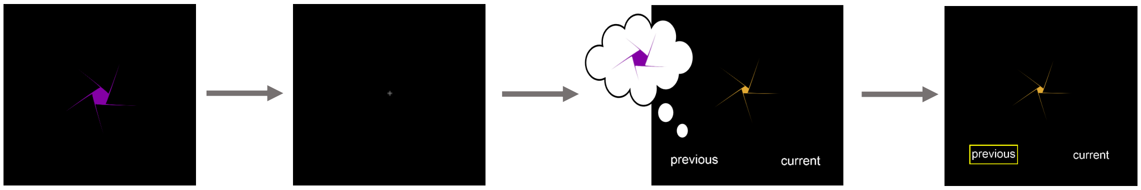


If you do not respond within 5 seconds you will see an **X** appear and the choice will be shown to you again at a later point. Please do your best to answer before the 5 seconds is up, but if you miss a shape or lose focus, it is better not to answer instead of making a random choice. Remember to try and choose a shape considering both the ***amount of $*** *and* ***chance of winning*** for each shape. You will complete three sessions today, lasting approximately 15–20 minutes each. At the end of *each session* one of your chosen shapes will be randomly selected and the associated 'gamble' will be played out. For example, if the selected shape is worth $5 and a 60% chance of winning, you will have a 60% chance of winning $5.
