## Extended Data for "A cognitive map of subjective value space for human risky choice"

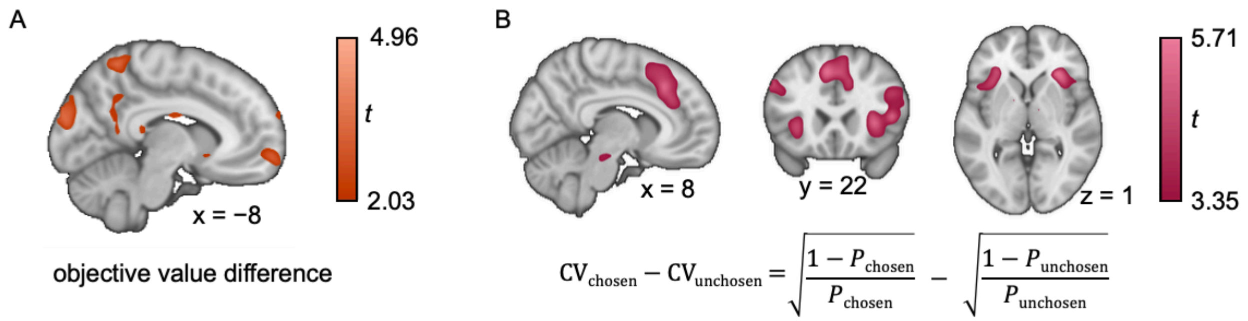


**Extended Data Figure 1. Neural effects objective value difference and risk.** We implemented the same GLM as with the subjective value difference analysis except calculated the objective value difference (**A**) and the coefficient of variation (CV; **B**)[1] of each option instead of the expected value. **A)** As with the subjective value difference map in the main text (Fig 1D), this map is thresholded at $P_{\text{uncorrected}} < 0.025$. **B)** We observe dMPFC/dACC, bilateral insula, and right lPFC significantly vary with the CV of the chosen relative to unchosen option (all $P\text{s}_{\text{TFCE}} < 0.05$). This demonstrates that our task recruits the same networks as other risky decision-making tasks (e.g., [2–4]). This map is thresholded at $P_{\text{uncorrected}} < 0.001$ for visualization.

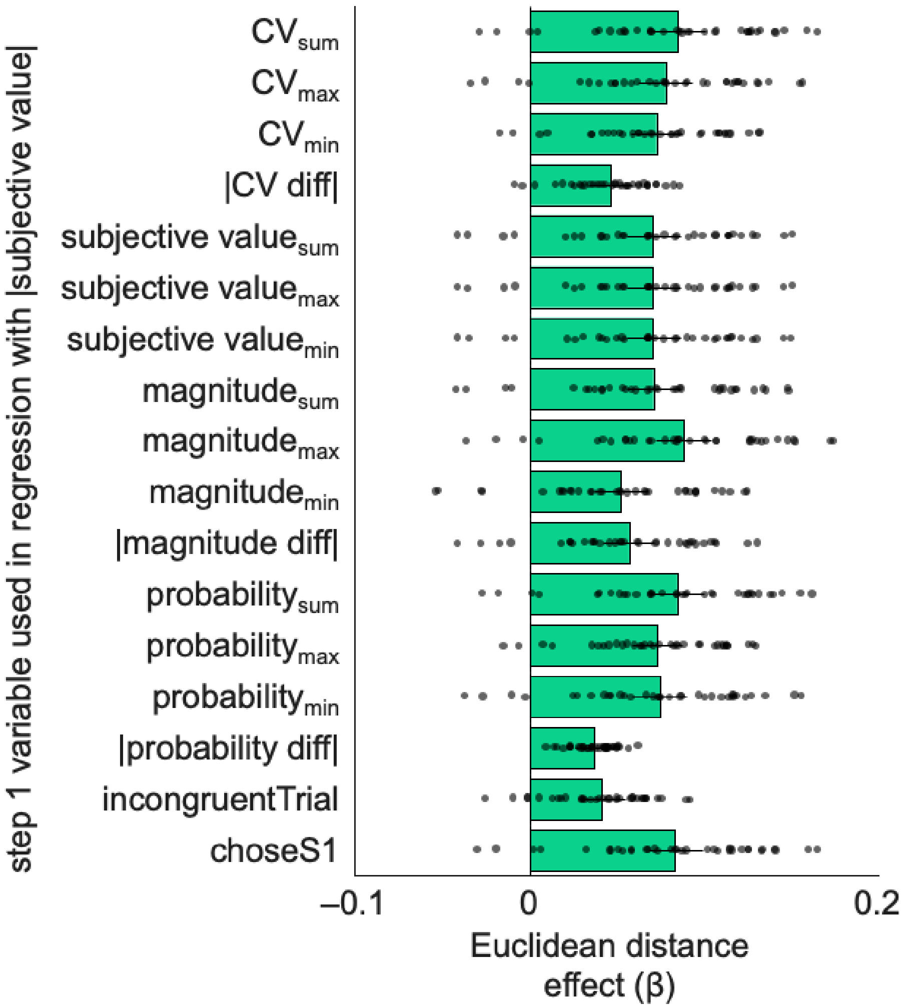


**Extended Data Figure 2. Euclidean distance between shapes in a decision vector predicts response time after accounting for other task-related variables.** To ensure the Euclidean distance effects are robust and not driven by other task-related variables, we implemented the hierarchical mixed effects regression procedure but in a series of additional regressions, adding a different additional variable at the first step for each regression. Each bar represents the Euclidean distance effect at the second level after accounting for the indicated variable at the first step of the regression. Across all variables, this effect remains significant (all *P*s $< 0.05$), and thus appears to be the primary driver of the observed effect. "choseS1" is a binary variable that indicates whether the first presented shape was chosen (0 if the second presented shape was chosen); "incongruentTrial" is a binary variable that indicates trials where probability was greater for one shape but magnitude was greater for the other shape (0 if that is not that case); "diff" indicates the difference between shapes; "min," "max," and "sum" subscripts refer to the minimum, maximum, and sum of the options for the indicated variable, respectively. Bar plots and error bars are the fixed effect and SEM of the estimate. The dots on are the reconstructed β values for each subject (i.e., the fixed effect β + random effects β for each subject).

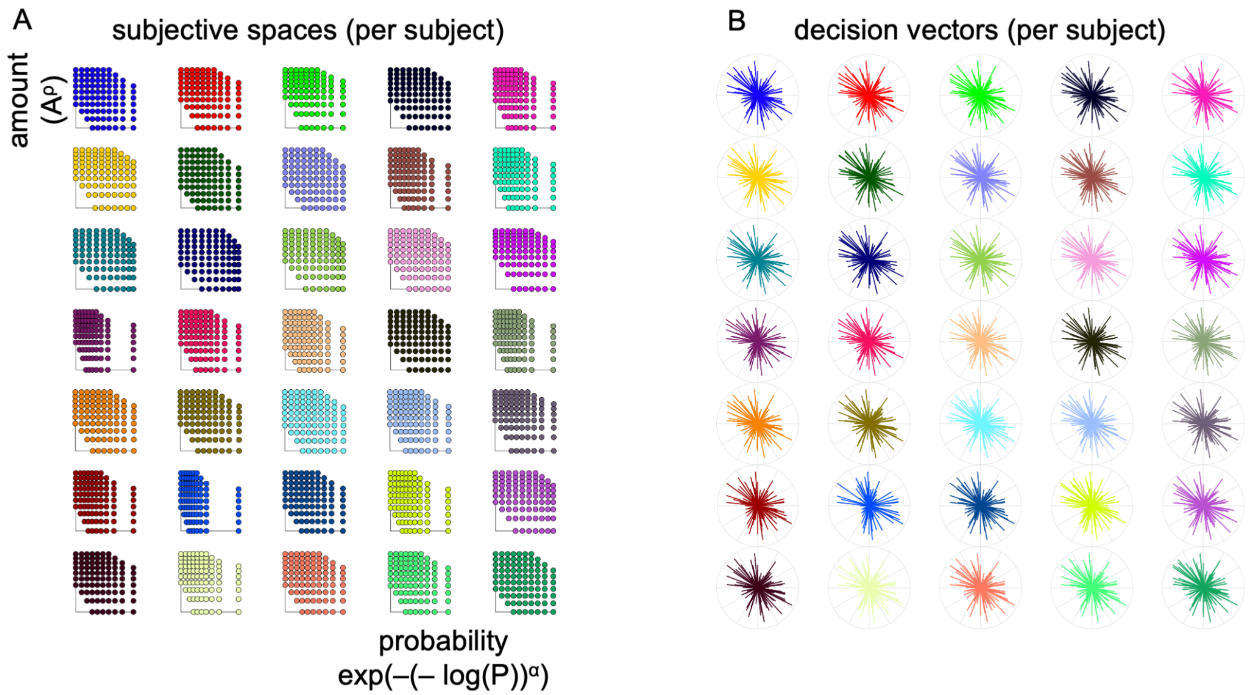


**Supplementary Figure 3. Subjective value spaces and decision vectors for all subjects. A)** The subjective value spaces are shown (as in Fig 2B, bottom left) for all subjects. **B)** The decision vectors in each space are shown on the right (as in Fig 2B, bottom right). Each subject is a unique color, and is located in the same place on the left and right plots. The subjective space and decision vectors for subject 5 (the example subject used in the main text) are in the top right.

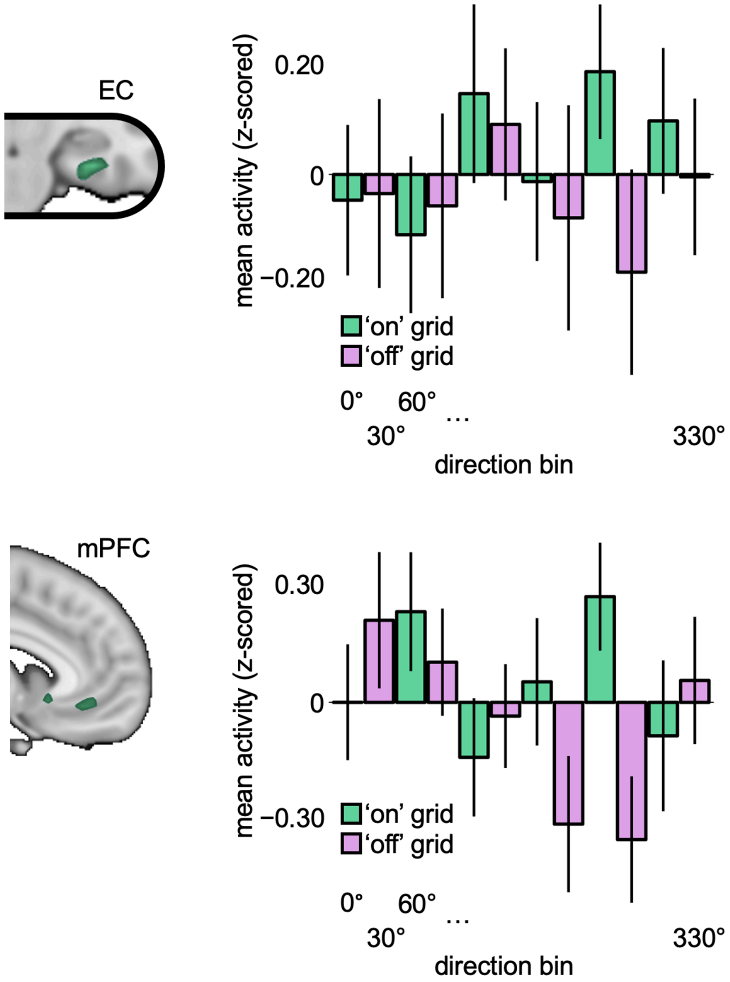


**Supplementary Figure 4. Grid-like representation is consistent across all decision vector directions.** To visualize the grid-like representation across the full 360º decision space, we estimated a third subjective $GLM_{grid}$ which included 12 parametric modulators (not orthogonalized). Each parametric modulator represented all choices in a given 30º bin (starting at −15º–15º). For each choice in a given bin, the value of this regressor was 1 (and was 0 for the rest of the bins), thus across all 12 bins all choices were represented. These bins represent the individual peaks and troughs of the cos(6θ) function where the odd numbered bins should have relatively high grid-like activity and the even numbered bins should have relatively low grid-like activity. This is shown for visualization purposes for the EC and mPFC voxels with $P_{uncorrected} < 0.05$. Mean activity (z-scored within participants) in EC (top) and mPFC (bottom) is shown separately for each 'on' (green) and 'off' (pink) grid bin based on the decision vector angles.

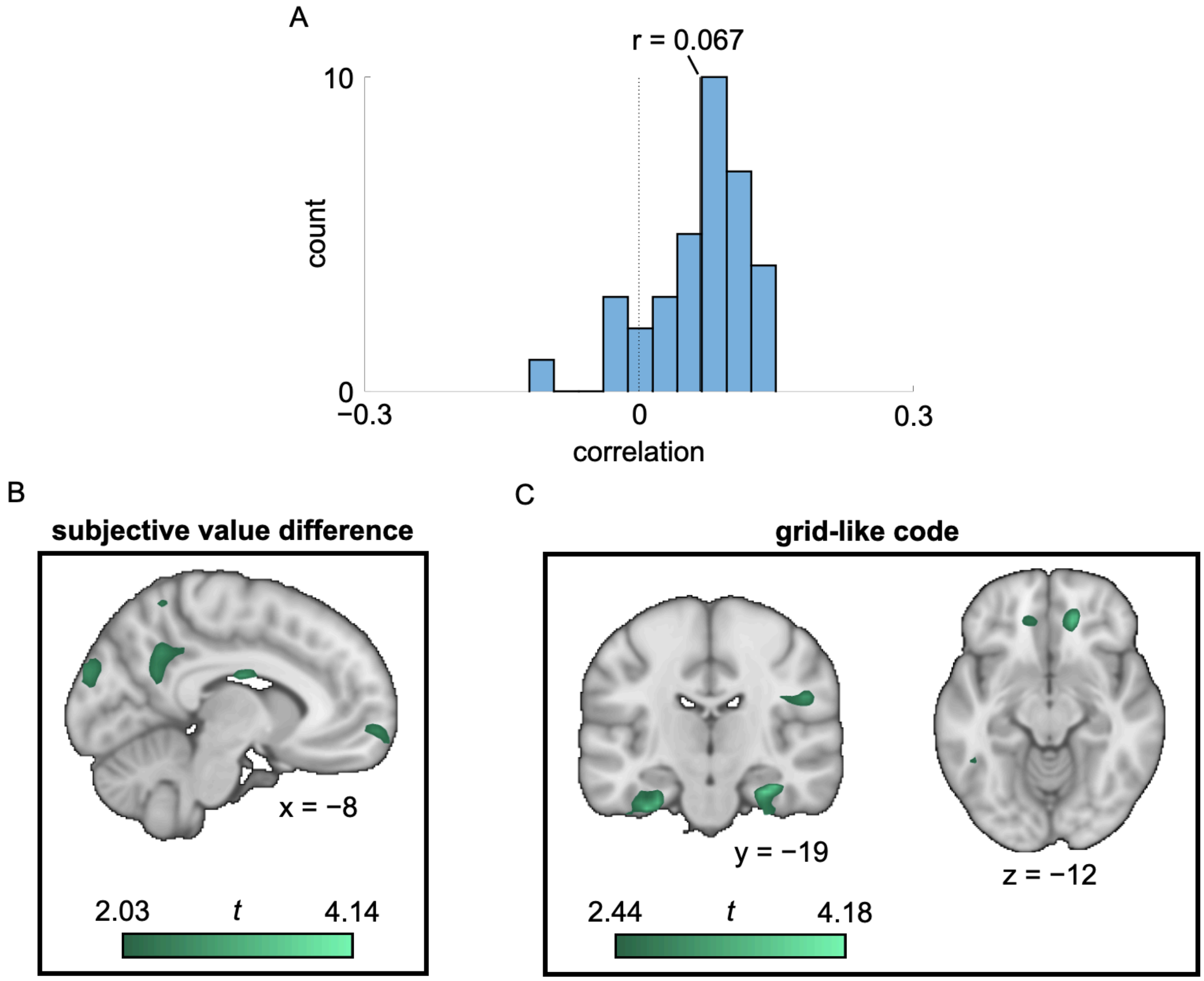


**Supplementary Figure 5. The subjective grid and subjective value results hold after controlling for correlations between regressors. A)** To confirm that the grid-like representation results aren't driven by a value signal, we first test if the subjective grid and subjective value regressors from the GLMs are correlated for each subject. The mean correlation was r = 0.067 (range −0.11 to 0.15). The distribution of r values across subjects is shown. Only three participants had significant correlations between the two regressors ($P < 0.05$; 8.6%). Because these regressors are uncorrelated across most subjects, and the number of participants with significant correlations is not different than would be expected from chance (chi-squared test vs 5%: $P = 0.33$), it is unlikely that the grid-like results can be explained by a value signal. **B, C)** Nonetheless, as another check, we additionally ran the same analyses from the main paper identifying a value signal (Figure 1D) and grid-like representation (Figure 3A) but included both regressors in the same GLM to allow them to compete for variance. **B)** We observed similar results as the primary analysis for subjective value difference (Figure 1D): significant rostral vmPFC ($P_{\mathrm{TFCE,\ SV}} < 0.05$) and marginally significant vPCC ($P_{\mathrm{TFCE,\ SV}} = 0.058$) positive parameter estimates and significant negative parameter estimates in bilateral anterior insula, bilateral dlPFC, and dACC (all $P\mathrm{s}_{\mathrm{TFCE}} < 0.05$). **C)** We also observed similar results as the primary analysis for grid-like representation of subjective value space (Figure 3A) where the EC and mPFC regressors remain significant (both $Ps_{TFCE,\ SV} < 0.05$). Finally, to ensure the correlations between second level estimates from Figure 5 aren't driven by a correlation between the first level regressors, we partialed out the correlation values for each subject from the correlation results. The results remain consistent with our original analysis showing that positive associations remain significant in PCC (vPCC: $\rho = 0.29$, $P = 0.050$; dPCC: $\rho = 0.37$, $P < 0.05$, partial rho; Figs. 5A and 5B) and significant or nearly significant in vmPFC (caudal vmPFC: $\rho = 0.37$, $P < 0.05$; rostral vmPFC: $\rho = 0.28$, $P = 0.055$, partial rho; Figs. 5C and 5D). Thus, our grid-like representation results are robust even after accounting for value coding (and vice-versa).

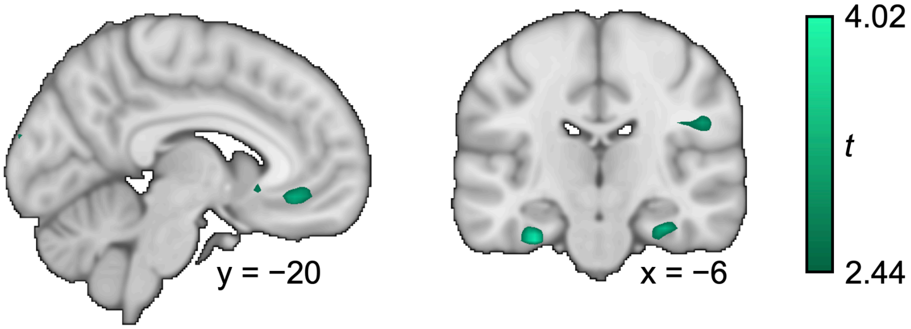


**Extended Data Figure 6. Grid-like effect for novel trials.** While less than 2% of trials contained choices participants previously saw during training, we nevertheless sought to confirm that the results are not driven by previously computed decision vectors. We tested for a subjective grid-like representation after excluding trials containing previously seen choices from first level GLMs. As with the original result, right ($P_{\text{TFCE, SVC}} < 0.05$), but not left ($P_{\text{TFCE, SVC}} = 0.16$) EC and mPFC ($P_{\text{TFCE, SVC}} < 0.05$) significantly encode a subjective grid-like code during novel choices. Thus, these data are consistent with the idea that cognitive maps are useful for generalization and novel choices.

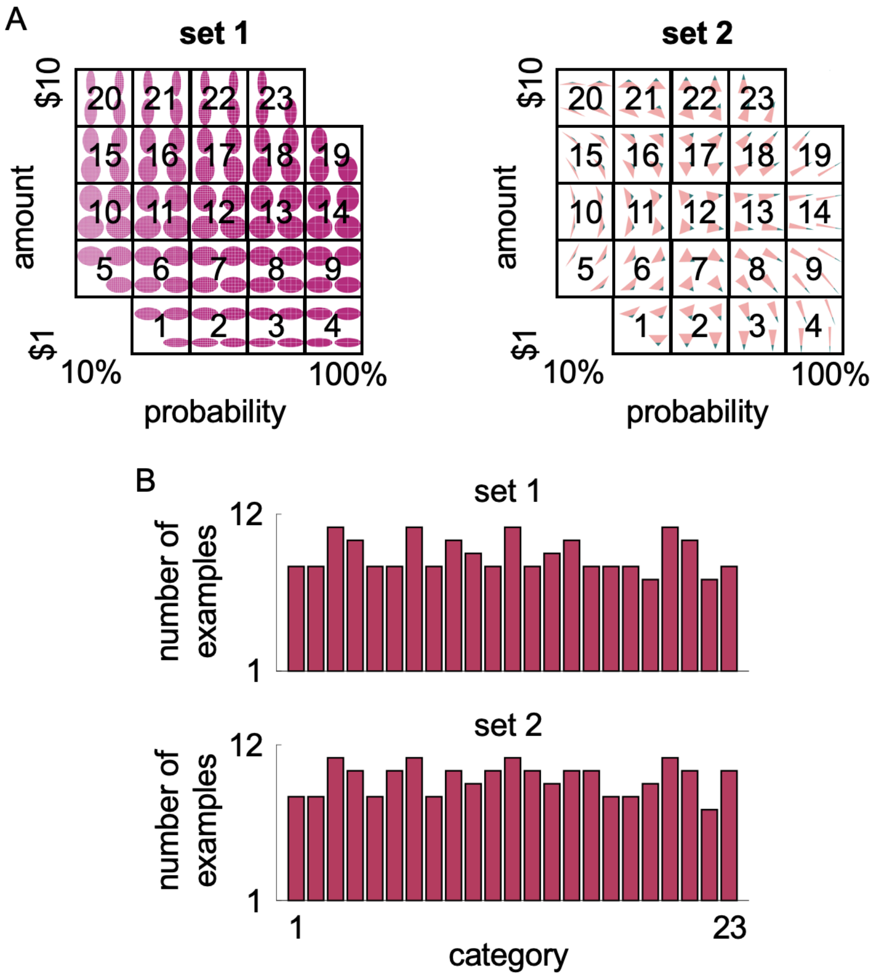


**Supplementary Figure 7. Categories used for the RSA. A)** To balance precision (i.e., location in the 2D space) and number of examples in each category, the shapes were placed into 23 groups of four as shown above. **B)** The number of examples per category was well distributed, as shown here for one example subject.

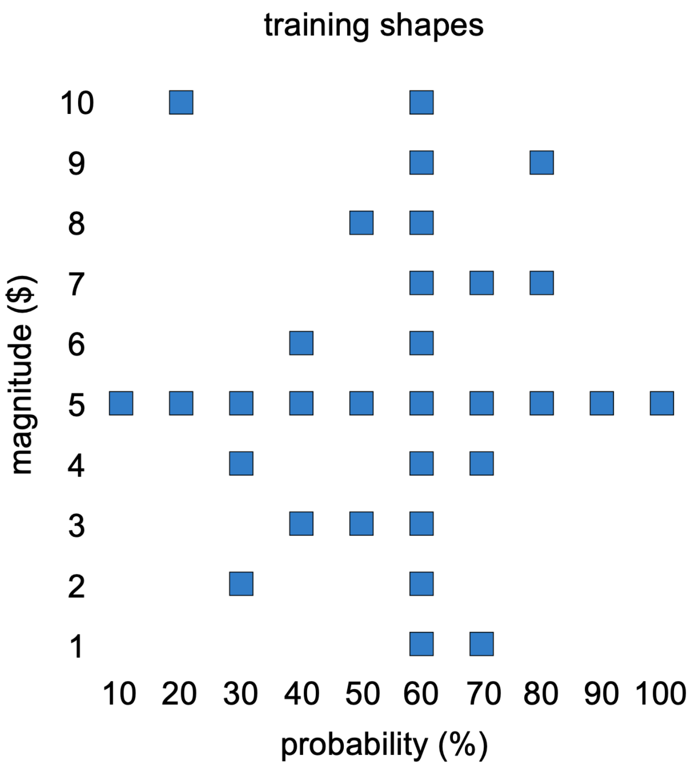


**Extended Data Figure 8. Shapes shown in training.** The location of the shapes used in training are shown by the blue squares. None of the other shapes were seen by the participants until they were in the scanner to participate in the task.
